# Prior low-severity fires reduce the risk of persistent forest loss from subsequent fires

**DOI:** 10.64898/2026.08.23.746551

**Authors:** Shengxi Gui, Shike Zhang, Yingtong Zhang, Jonathan A Wang, Zhe Zhu, Thiago Gonçalves-Souza, Mohammed Ombadi, Yanlan Liu, Jin Tang, Peter B Reich, Benjamin P. Goldstein, Kai Zhu

## Abstract

Intensifying fire regimes threaten forests globally^1,2^, but the risk of persistent post-fire forest loss^3,4^ and its potential mitigation remains poorly quantified. We analyzed millions of wildfires worldwide from 2001 to 2024 and tracked recovery in satellite-observed forest structure and ecosystem function^5,6^. Post-fire persistent forest loss, indicated by modeled non-recovery to pre-fire conditions over decadal timescales, affected 57.1% ± 1.4% (mean ± s.e.) of burned forest area globally since 2001, with hotspots in Pacific temperate and southern boreal forests. We then identified “crucial fires” as events exceeding a stringent modeled-risk probability threshold for persistent structural or functional non-recovery, with fire severity strongly predicting this loss. This severity dependence revealed a management pathway, as locations with prior low-severity fire experienced lower severity in subsequent wildfires^7–9^ and had lower modeled probability of becoming crucial. Under a model-based counterfactual scenario, applying the estimated severity attenuation was associated with a 7.6 ± 0.6% reduction; the top 1% of road-accessible areas accounted for 35% of this reduction. These results provide a global framework for identifying where wildfire threatens forest resistance and where targeted low-severity fire management like prescribed fire^8,10^ might be used to combat global forest loss.

## Main

As climate extremes intensify and reshape fire regimes, forests are increasingly exposed to fires capable of causing long-term ecological change^1,11,12^. Disturbance, including wildfire, is a natural component of many forest ecosystems, and recovery toward pre-disturbance conditions is often expected. Yet burned area alone does not determine ecological impact. Some burned forests recover to their pre-fire structure and function, but a growing number of such areas remain degraded for decades or transition toward alternative landscapes^3,13^. Because forests regulate the Earth system through carbon storage and biophysical exchanges of energy and water, persistent failure to recover can reduce the long-term contribution of affected landscapes to the global forest carbon sink^14,15^. A key challenge is distinguishing forest fires followed by recovery, including delayed regeneration, from fires that push forests beyond their recovery capacity^5,16,17^. Resolving this challenge is needed to quantify the ecological costs of contemporary fire regimes and for identifying effective strategies to reduce persistent post-fire forest loss.

Post-fire forest recovery is a multidimensional process that involves both rebuilding forest structure and restoring ecosystem function^6^. Large-scale assessments often rely on greenness-based indicators such as the Normalized Difference Vegetation Index (NDVI)^18^, which can detect vegetation return but may not distinguish forest recovery from replacement by grasses, shrubs, or sparse woody cover^6,19^. A post-fire landscape may therefore appear green while still failing to regain its former canopy structure or biogeochemical cycles^3,4^, allowing apparent regrowth to mask degradation. We therefore separate recovery into two complementary dimensions^5,6^: forest structure, describing canopy spatial and vertical coverage, and ecosystem function, reflecting carbon and water exchange with the atmosphere. Because these dimensions can recover at different rates and toward different end statuses, apparent recovery in one dimension may mask persistent change in another, indicating post-disturbance reorganization rather than full resilience to fire. In this study, post-fire forest resilience is assessed by the failure for burned forests to regain pre-fire structural and functional conditions. Tracking both dimensions helps distinguish apparent regrowth from failure to recover.

Despite its global importance, persistent post-fire non-recovery of forest structure or ecosystem function is rarely evaluated as a characteristic of individual fire events, even though it represents a significant long-term ecological consequence. At the event scale, wildfires are commonly described by occurrence^12^, extent^20^, and immediate severity^21^, yet similar events can follow divergent recovery trajectories depending on vegetation and environmental context^22^. Persistent non-recovery can therefore have lasting consequences for carbon storage^23^, hydrological function^24^, and biodiversity^25^. We define fires with high probability of persistent non-recovery as “crucial fires”. This concept seeks to link wildfire impacts to tipping-point perspectives on forest resilience by treating modeled long-term non-recovery as an event-level consequence, without equating it with a permanent regime shift^26–29^. It also remains unclear how widespread these high-risk fire events are globally, whether they are becoming more frequent, or whether their distribution is shifting among regions. To our knowledge, no current study maps this ecological consequence perspective to the individual event or evaluates how it relates to fire attributes and environment. Fire size may influence recovery by determining the spatial extent of seed-source loss and landscape fragmentation, whereas fire severity determines the immediate magnitude of canopy, biomass, and functional damage^22,30,31^. Quantifying how these attributes, combined with environmental factors like temperature and precipitation, can provide key insights into which fires are most likely to cause persistent forest loss and which landscapes are most at risk.

Among event-level fire attributes, severity is especially relevant because it can be influenced, at least in part, by fuel structure, stand condition, and prior disturbance history^7,10^. This dual role raises a management question: do prior fire regimes influence subsequent wildfire severity and thereby lower the risk of persistent forest non-recovery? Prescribed fire is typically the intentional use of low-severity fire to reduce fuels, and prior low-severity wildfire can provide a natural analog^32^. Regional studies on western U.S. suggest that low-intensity or low-severity fire reduces the risk of subsequent high-intensity fire^8,33^ and offsets climate-driven declines in regeneration^22^. Low-severity fires that consume surface and understory fuels, decrease vertical and horizontal fuel continuity, and modify stand structure may reduce the severity of subsequent wildfires^7,34^. If so, prior low-severity fire could attenuate future crucial fire risk, and suggest the latter as an effective strategy to mitigate^10^. However, the link between prior and subsequent wildfire severity remains unclear over time and across landscapes.

Here, we examine around 3.1 million individual fire events to map persistent losses globally from 2001 to 2024 and clarify the link between risk of crucial fire and historical fire severity. We combine global wildfire records with satellite-based recovery trajectories of forest structure, represented by tree cover (TC) and leaf area index (LAI), and trajectories of ecosystem function, represented by net primary productivity (NPP) and evapotranspiration (ET)^35,36^(**Methods**). After excluding intentional fires to clear land, we quantify where burned forests fail to regain pre-fire structure or function. By training spatially cross-validated XGBoost models on these recovery outcomes (**Methods**), we estimate event-level probabilities of persistent non-recovery from fire size, fire severity, and environmental conditions to classify crucial fires, map their global distribution, and identify key risk drivers. Finally, we test whether prior low-severity fires, as a natural analog for prescribed fire, reduce crucial fire risk, and identify road accessible areas where low-severity burning is both viable and beneficial. By moving from wildfire consequence to mechanism and mitigation, our analysis provides a global framework to identify where wildfire threatens forest resilience and where targeted forest management should be prioritized to effectively reduce persistent forest loss.

## Results

### Persistent post-fire forest loss is widespread, heterogeneous, and multi-dimensional

Across global fire events from 2001 to 2014 and tracked through 2024, post-fire recovery was limited and highly heterogeneous across space. To illustrate the basis of our trajectory-based identification, the paired case study in **Fig. 1A-B** indicates the global pattern based on modeled recovery potential: even nearby burned forest patches can follow markedly different trajectories, with some remaining far below their pre-fire baseline for nearly two decades and projected to remain below it, while others recover toward it within 13 years. Scaling this trajectory analysis globally revealed widespread persistent forest loss, defined here as cases in which the modeled long-term recovery asymptote remained below the pre-fire state. Persistent loss was defined by the strongest modeled asymptotic deficit across tree cover, LAI, NPP, and ET (**Methods**). Independent land-use filters minimized contamination from agricultural fires and post-fire conversion, so these estimates primarily reflect post-fire recovery in the natural environment (**Fig. S1**). Globally, across fires from 2001-2014, 19,215 ± 419 km² (mean±s.e.) of burned forest was classified as persistently non-recovering based on the modeled long-term asymptote per year. Indicator-specific persistent non-recovery was most pronounced for tree cover, NPP, and ET, affecting 42.3% ± 1.8%, 47.9% ± 2.6%, and 45.0% ± 1.2% of burned forest area, respectively, and was lower but still widespread for LAI (29.0% ± 1.9%). After accounting for overlap, 57.1% showed non-recovery in at least one indicator. Persistent losses in both forest structure and ecosystem function were highly concentrated in specific regions, including Pacific North America, central Africa, mainland Southeast Asia, and Siberia (**Fig. 1C**). Across the global forest biome, these impacts exhibited strong spatial heterogeneity in both loss density and recovery proportions (**Extended Data Fig. 1**). Independent U.S. Forest Inventory and Analysis (FIA)^37^ observations across nine ecoregions yielded recovery proportions broadly comparable to MODIS-based structural estimates from forest structure (tree cover and LAI) (nRMSE = 0.34; **Text S4**, **Fig. S18-20**).

**Figure 1.**
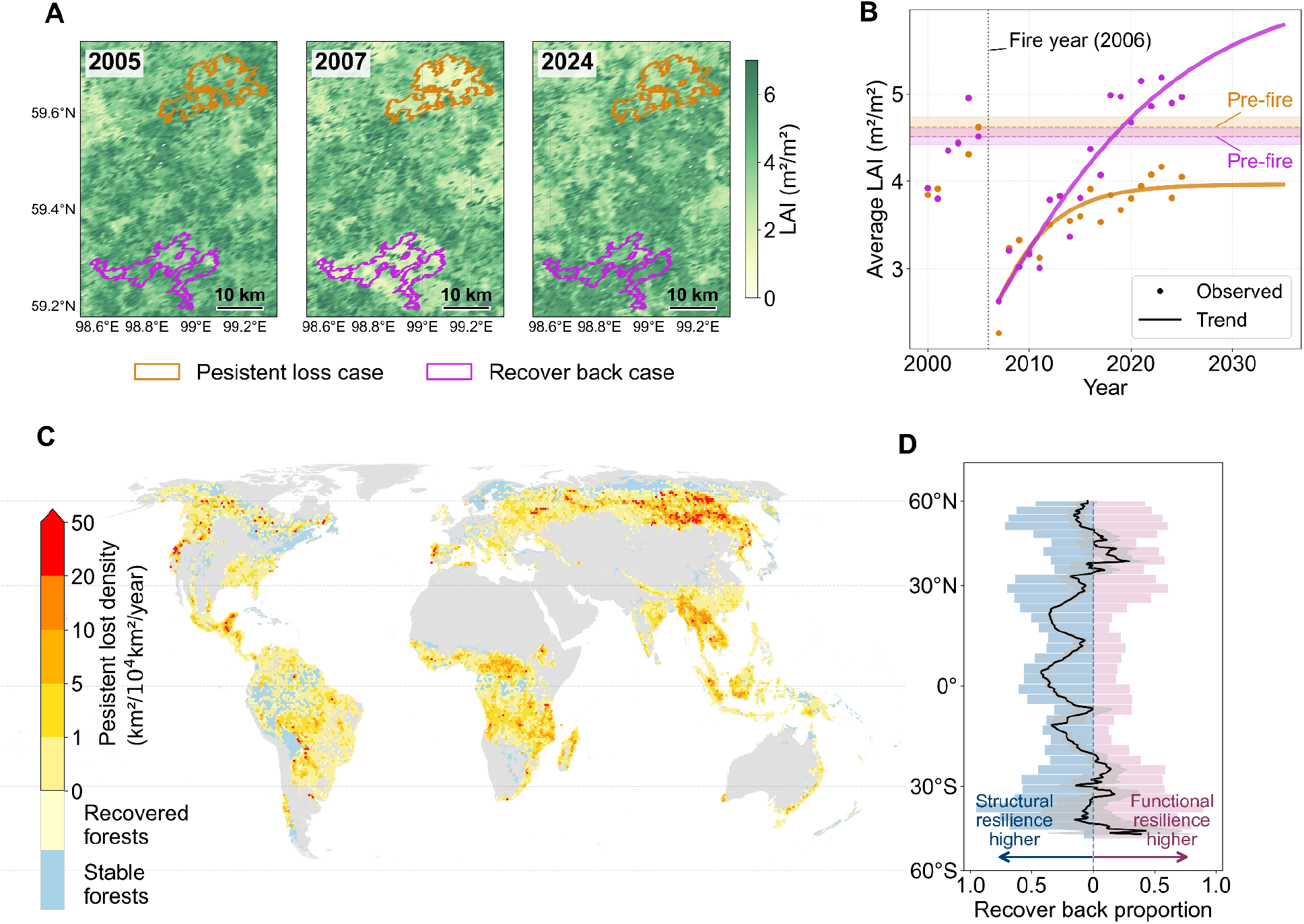
Contrasting local trajectories and global patterns of postfire forest resilience. (A) LAI maps for a boreal case study in Siberia for the fire year 2006, showing before fire (2005), immediately after fire (2007), and in 2024, with one case of persistent loss and one that recovered toward pre-fire conditions. (B) LAI time series for the two outlined cases, with points showing observations, colored curves showing fitted recovery trajectories, horizontal dashed lines marking prefire LAI, and the vertical dotted line marking the 2006 fire year. (C) Global density of persistent forest loss summarized on 50-km equal-area hexagons (fires from 2001-2014); warm colors indicate higher persistent-loss density, whereas pale yellow and blue denote recovered forests and stable forests (without wildfires or postfire vegetation indicators are the same or higher than prefire condition), respectively. Values were derived from the maximum signal across structural (TC, LAI) and functional (NPP, ET) indicators. (D) Latitudinal profile of resilience contrast between structural and functional dimensions; the black curve shows the mean contrast with uncertainty shading, and the side histograms show the corresponding latitudinal distributions for structural and functional resilience.

Beyond its spatial extent, persistent forest loss also differed in the structural and functional dimensions of recovery. Their contrast was modest overall but varied with latitude: the two dimensions were broadly similar at high northern latitudes, whereas structural recovery more consistently exceeded functional recovery between tropical regions (**Fig. 1D**). In temperate and boreal forests, rapidly growing early-successional vegetation may sustain productivity even when tree regeneration and canopy recovery are delayed or incomplete^38,39^. At lower latitudes, canopy cover or leaf area may recover before carbon uptake and water cycling because young or compositionally altered vegetation remains functionally distinct from the pre-fire forest^40^. Conversely, where functional recovery exceeded structural recovery, rapidly growing early-successional vegetation may restore productivity and water flux before mature canopy structure is re-established. These contrasting patterns show that burned forests may fail to recover in different ways across biomes, and that post-fire forest loss is fundamentally multidimensional.

### Crucial fires show regional heterogeneity and are distinguished by severity

Moving from spatial patterns of persistent-loss to individual fire events, crucial fires showed contrasting regional trajectories without a clear global trend. We defined a crucial fire as an event with less than 25% predicted recovery probability in any structural or functional indicator (**Methods**), representing a conservative threshold for persistent loss in at least one recovery dimension. The main patterns were robust to alternative thresholds (**Text S3**; **Fig. S11**). The annual forest area burned by crucial fires did not exhibit a consistent global increase or decrease from 2002 to 2023 (**Fig. S3**). The difference in trend from all fire events and slight increase (+0.03%/yr) in the crucial fire proportion suggests that changes in crucial fire activity were not solely driven by changes in overall fire extent (**Extended Data Fig. 2A-C**). Spatial and biome-level^41^ patterns revealed contrasting (**Fig. 2A-B** and **Fig. S4**). Crucial burn area declined in parts of tropical moist broadleaf forests, including southeastern Amazonia, but increased across African savanna and savanna–forest regions. The largest biome-level increase occurred in forest-fire area within the tropical grasslands and savannas biome. Boreal and temperate coniferous forests showed persistent activity and weak or non-significant increases, whereas several smaller biomes were dominated by stable or low-magnitude trend classes. Therefore, recent changes in crucial fire activity reflected spatial heterogeneity rather than uniform global expansion.

**Figure 2.**
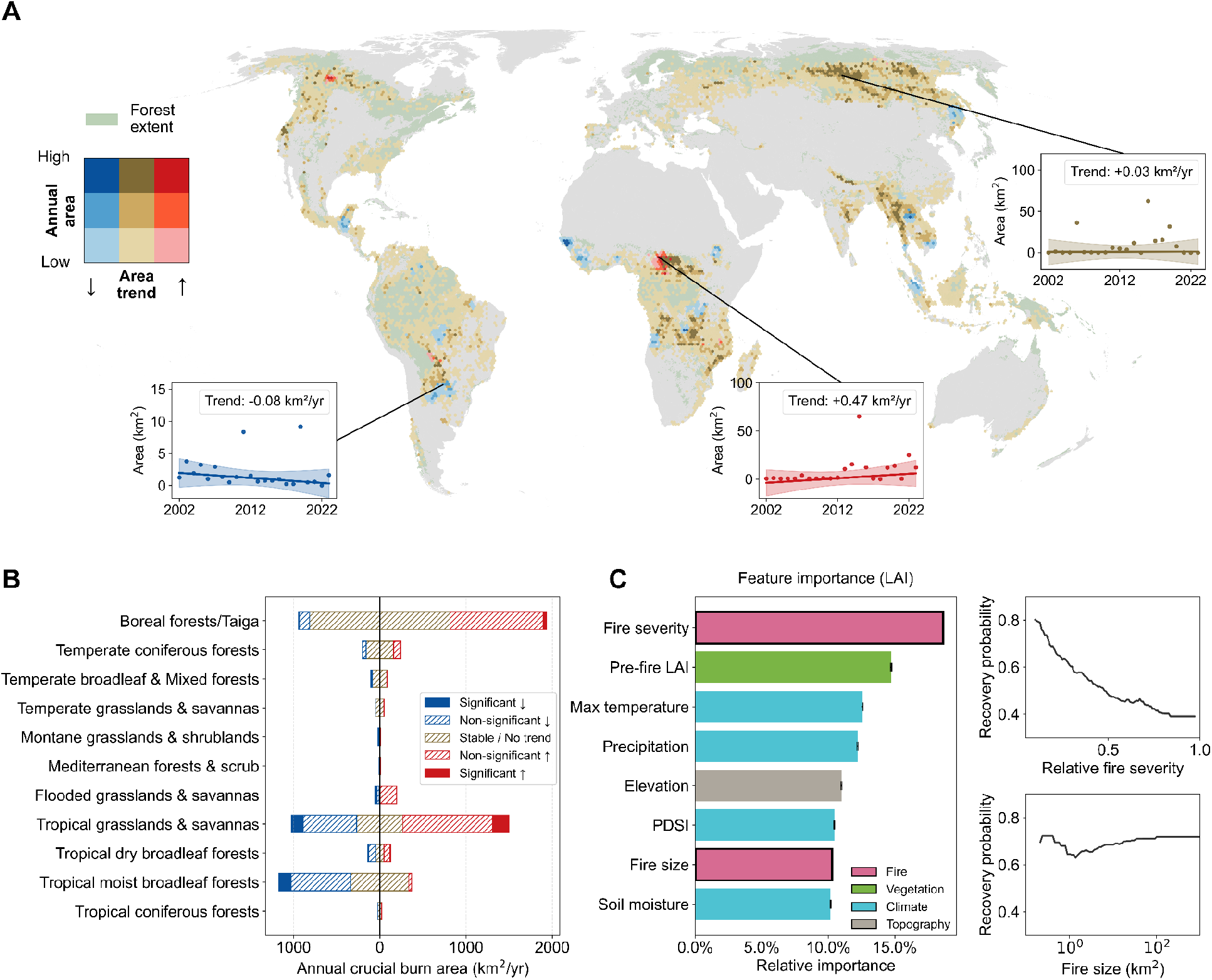
Spatial trends and drivers of crucial fires. (A) Global map of crucial fire regimes summarized on 50-km equal-area hexagons, combining mean annual crucial fire burned forest area and temporal trend in annual crucial fire area. Blue tones indicate decreasing area trends (<-0.01 km^2^ / yr), red tones indicate increasing area trends (> 0.01 km^2^ / yr), and darker colors indicate larger mean annual crucial fire area. Insets show sample local time series from regions with contrasting trajectories. Forest extent is shown in green. (B) Biome-level summary of annual crucial fire area with their trend, partitioned into significant decrease, non-significant decrease, stable/no trend, non-significant increase, and significant increase classes. Non-forest dominated biomes may still contain forest patches, and only burned areas within those forest pixels were included. (C) For the LAI-based recovery model (XGBoost), feature-importance scores identify fire severity, pre-fire LAI, and climate variables as the main predictors of recovery probability; partial dependence plots show declining recovery probability with increasing fire severity and relatively stable probability with fire size change.

Classification models showed that crucial fires were distinguished more by severity than size (**Methods**). In the LAI-based recovery model as representative, fire severity was the strongest predictor of recovery probability, with pre-fire vegetation and climate variables all more important than fire size (**Fig. 2C** and other metrics in **Fig. S2**). Partial dependence analysis showed that predicted recovery probability declined by approximately 30% with increasing relative fire severity but varied markedly less 10% across fire size (**Fig. 2C**), with only weak correlations between the two attributes across indicators (**Fig. S16**). This pattern was consistent across recovery dimensions: severity ranked first in the LAI model and second in the tree-cover and ET model, whereas climate and pre-fire conditions also contributed strongly to functional recovery, with severity exceeding fire size in the NPP model. After accounting for vegetation and environmental conditions, crucial fires were distinguished more by high severity than by large fire size.

### Low-severity fire legacies reduce subsequent severity and crucial fire risk

Because low-severity fires were more likely to remain within forest recovery capacity, we tested whether their legacies also reduced subsequent wildfire severity and thereby lowered the probability of persistent non-recovery. For reburned locations, we matched pixels that had burned previously at low severity to nearby pixels in the same subsequent wildfire with similar pre-fire vegetation but no prior-burn history, thereby reducing differences in event-level fire weather and other conditions (**Methods**; **Extended Data Fig. 3**; **Fig. S5**). A complementary analysis found no strong or consistent association between subsequent severity and monthly fire-weather indicators (**Text S5** and **Fig. S17**). Within these matched pairs, areas with a previous low-severity burn experienced lower severity in the subsequent wildfire than areas without a previous burn (**Extended Data Fig. 4**). This effect indicates greater resistance to severe reburning, not necessarily greater resilience after repeated fire, because short-interval or high-severity reburns can still impede regeneration and promote persistent forest loss^4,42^. Across both structure and function levels (TC, LAI, NPP, and ET), low-severity prior-burn pixels generally showed lower subsequent fire severity than matched no-prior-burn pixels with a mitigation effect, and this effect was strongest under low-to-intermediate pre-fire vegetation conditions (**Fig. 3A** and **Fig. S6**). However, higher-severity prior fire showed no consistent protective effect and sometimes increased subsequent severity with an exacerbation effect, indicating that mitigation was isolated to low-severity fire legacies rather than prior burning in general (**Fig. 3B**, **Extended Data Fig. 4** and **Fig. S6-7**). This contrast remained robust after accounting for uncertainty in the estimated effects (**Fig. S29**).

**Figure 3.**
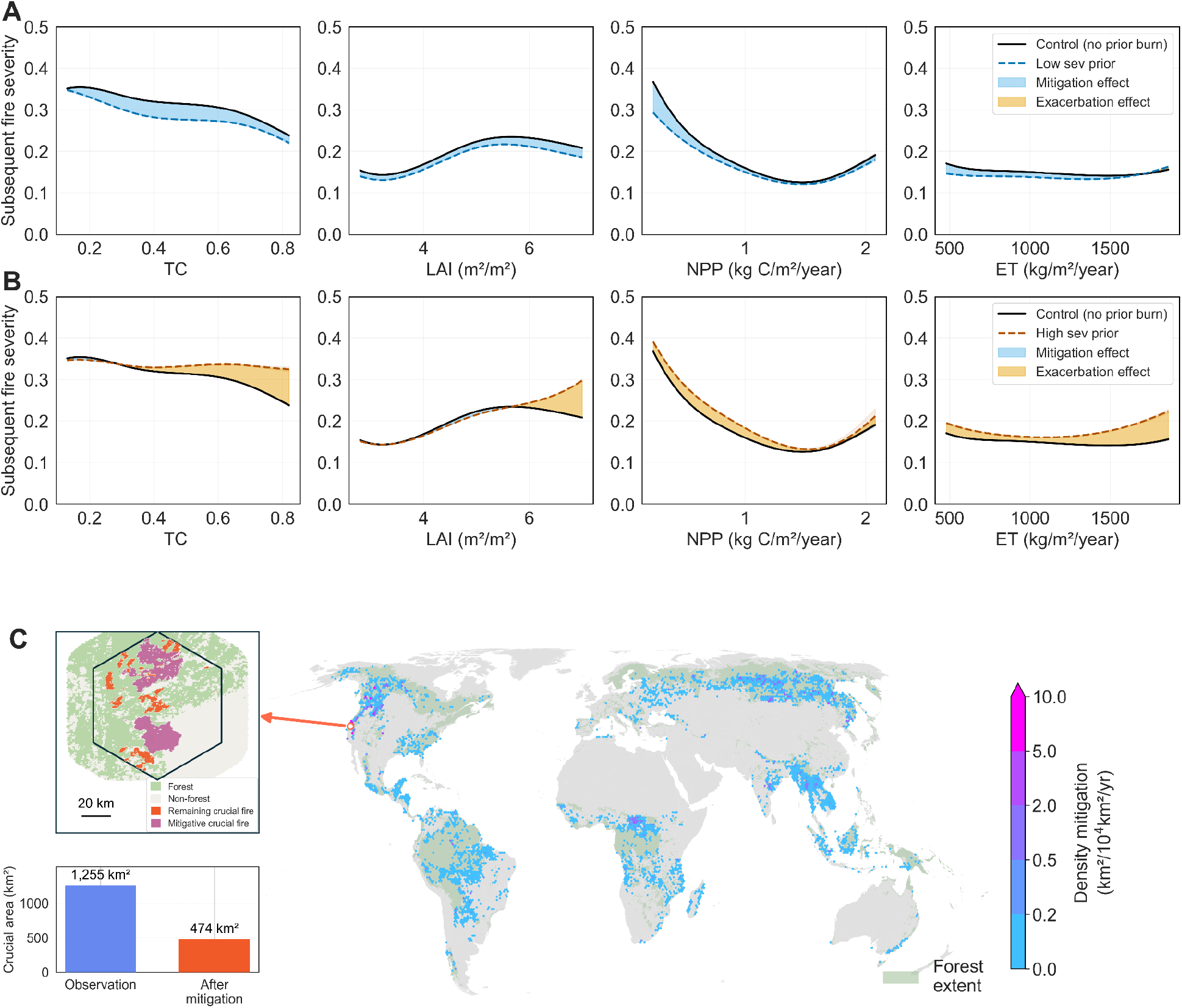
Low-severity prior burning reduces subsequent fire severity and the crucial fire area. (A) Relationships between subsequent fire severity and pre-fire vegetation condition under low-severity prior burning, compared with matched control areas without prior burn. (B) Corresponding relationships under high-severity prior burning. Columns show structural indicators (TC and LAI) and functional indicators (NPP and ET). Solid black curves indicate controls, dashed colored curves indicate prior-burn conditions, and shaded areas indicate mitigation or exacerbation effects. Blue shading indicates lower subsequent fire severity relative to controls, whereas yellow shading indicates higher subsequent fire severity. (C) Global reduction in annual crucial fire density associated with low-severity prior burning, summarized on 50-km hexagons and averaged across the four vegetation indicators. Colors indicate annual reduction in crucial fire area normalized by hexagon area, and green shading shows forest extent. The inset shows one highlighted 50-km hexagon in a temperate coniferous forest landscape in western North America, with forest, non-forest, mitigated crucial fire area, and remaining crucial fire area. The bar chart compares the modeled crucial fire area within this hexagon under observed conditions and under the low-severity prior-fire scenario.

The severity attenuation associated with prior low-severity fire translates into a lower risk for crucial fires at the global scale. We then applied this severity reduction in the recovery models to estimate how many fires would no longer be classified as crucial under the low-severity prior-fire scenario. To translate this severity reduction into global mitigation potential, we reduced each fire’s observed severity by the corresponding effect estimated from the matched prior-fire comparisons, held all other predictors constant, recalculated indicator-specific recovery probabilities, and reapplied the same 25% classification threshold. Relative to the observed-severity baseline, this counterfactual reduced the modeled forest area burned by crucial fires by 413.7 km²/yr, or 7.6%. The global mitigation effects were spatially uneven, with stronger effects concentrated in western North America, boreal forest zones, and savanna-forest mosaics (**Fig. 3C**, and uncertainty map **Extended Data Fig. 5**). In one highlighted temperate coniferous hexagon in western North America, the modeled crucial fire area declined from 1255 km^2^ under observed conditions to 474 km^2^ after the mitigation scenario (**Fig. 3C**). This local example illustrates how reducing subsequent fire severity can shift individual fire events above the recovery-probability threshold and out of the crucial fire category. Biomes also differed in absolute and fractional mitigation potential: extensive fire-prone biomes contributed large absolute reductions, whereas temperate coniferous forests and Mediterranean forests and scrub showed the largest fractional reductions (**Extended Data Fig. 6**). These results suggest that low-severity fire legacies can reduce modeled crucial fire risk where they substantially lower subsequent fire severity.

### Mitigation benefits are concentrated in accessible high-benefit areas

We find that mitigation opportunities are highly uneven across global forests. For each road-accessible 50-km grid cell, we calculated the mitigation area as the forest burned area classified as crucial under observed conditions but no longer classified as crucial under the low-severity prior-fire scenario. We then ranked road accessible grid cells by mitigation area to find highly beneficial regions. The resulting high-benefit areas were strongly clustered (**Fig. 4A** and **Fig. S8**), and this concentration persisted through time (**Fig. 4B**). The top 1% of road accessible landscapes accounted for 35% of total modeled mitigation potential, and the top 10% captured nearly 73%; the remaining 90% contributed only about 27% (**Fig. 4C**). Thus, targeting a small fraction of areas could capture disproportionately large benefits from prescribed fire and other severity reduction management.

**Figure 4.**
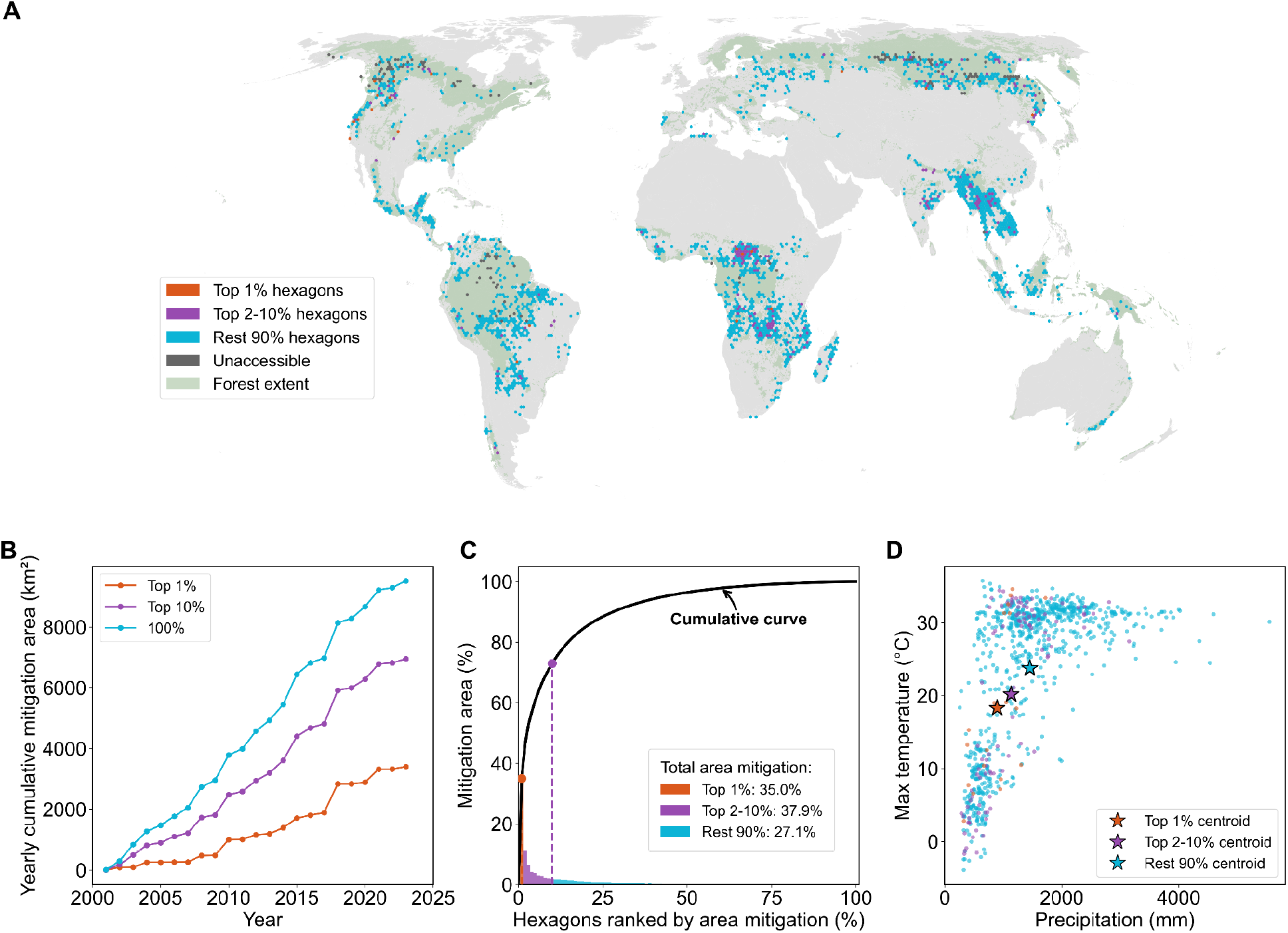
Accessible priority landscapes capture a disproportionate share of potential mitigation benefits. (A) Global distribution of treatment-priority classes on the 50-km equal-area hexagon grid. Hexagons were ranked by realized mitigation area among road-accessible locations, with colors indicating the top 1%, top 2 to 10%, remaining 90%, and inaccessible hexagons (identified using road-network accessibility); green shading denotes forest extent. (B) Accumulated mitigation area through time under three treatment scenarios: treating all accessible hexagons, only the top 10% of ranked hexagons, and only the top 1%. (C) Distribution of hexagon contributions to the total mitigation area and the corresponding cumulative saved-area curve after ranking hexagons from highest to lowest mitigation area. (D) Climate space of priority classes for hexagons with >5% mitigation of crucial burned area, showing mean precipitation and maximum temperature, with stars marking class centroids. The mitigation area was defined as the forest-burned area of fires classified as crucial under observed conditions but not under the counterfactual low-severity prior-burn scenario, aggregated across the TC, LAI, NPP, and ET models.

The highest-benefit accessible areas also occupied a distinct climate space, tending to occur in hotter and drier conditions. Among landscapes with substantial mitigation potential, the highest-ranked areas tended to be colder and drier than low-priority accessible landscapes (**Fig. 4D**). Together, these results suggest that the opportunity to reduce crucial fire area is not globally diffused. Instead, a relatively small number of accessible, climatically distinctive landscapes dominate the modeled mitigation potential, suggesting that low-severity fire management could be most effective when spatially targeted.

## Discussion

We provide a global, event-level assessment of wildfire consequences by linking individual forest-fire events to long-term trajectories of structural and functional recovery. This approach identifies fires with high modeled probability of persistent post-fire non-recovery and evaluates where low-severity fire legacies may reduce that risk. Although wildfires are commonly assessed through occurrence, burn area, or immediate severity^43–45^, our findings show that their long-term ecological consequences follow distinct spatial and temporal patterns and cannot be inferred from fire extent alone.Previous fire-management studies have shown that prescribed or low-severity fire can reduce subsequent wildfire severity^8–10,32^, our global analysis extends this evidence by showing that the associated severity reduction may also lower the modeled probability of persistent forest non-recovery. More broadly, our framework provides a global assessment for prescribed or managed fire by identifying landscapes where low-severity burning is ecologically plausible, operationally accessible, and most likely to reduce future crucial fire risk.

The concept of crucial fires provides an event-level diagnostic of wildfire consequence by identifying a recovery-defined subset of mapped forest-fire events with high modeled probability of persistent non-recovery, rather than ranking fires by extent alone. In our analysis, these low recovery probability fires represented a minority of mapped forest-fire events but captured a distinct dimension of wildfire impact: the risk that burned forests may not quickly (or ever) return to their pre-fire structural or functional condition. Despite the challenges of long-term recovery prediction^46^, integrating fire severity with pre-fire vegetation, climate, and terrain allows our framework to harness early post-fire observations to provide a screening-level early-warning estimate of persistent non-recovery risk. Through this recovery-threshold lens, global fire change appears less as a uniform increase than as a spatial redistribution of high-consequential fire events across regions and biomes (**Fig S4-S5**), diverging from broader fire-activity attributes such as declining global burned area^47,48^ and rising extratropical forest fires^49^. Across all model-valid forest fires, total burned area declined slightly, whereas the proportion classified as crucial increased, indicating that crucial fire trends were not driven by overall fire extent alone (**Extended Data Fig. 2**). In the southeastern Amazon, however, declines in crucial fire area paralleled declines in total forest-fire area after strict filtering of likely land-use fires, suggesting that reduced overall fire exposure, rather than a broad reduction in recovery risk, was the cause^50,51^. In contrast, increases in African savanna-forest transition zones point to rising risk in flammable land-use mosaics^47,52,53^, while boreal patterns remain episodic and dominated by extreme fire years^12,54^.

Low-severity fire legacies were associated with lower subsequent fire severity and lower modeled crucial fire risk, but this effect should be interpreted as a targeted, context-dependent pathway rather than a universal fire-management solution. In most biomes, prior low-severity fire lowered subsequent severity (**Fig. S6**-**S7**), reducing the probability of future crucial fires^7,10,11^. This protective effect is consistent with fuel-mediated mechanisms documented in regional studies^9,55,56^. Low-severity fire can consume surface and ladder fuels while retaining live overstory structure^8,34^, thereby reducing fuel continuity and helping counteract fuel buildup associated with prolonged fire suppression in many western U.S. forests^11,57^. In contrast, high-severity fire can remove canopy cover, increase deadwood, dry the surface environment, and promote understory fuels^56,58,59^ that may favor more severe reburning than in adjacent forests (**Fig. S12**). The contrast between low-and high-severity prior burns therefore suggests that mitigation depends not on prior burning alone, but on whether prior fire creates fuel and stand conditions that reduce subsequent severity without generating high-severity legacies.

The modeled mitigation effect was relatively modest globally but highly concentrated spatially, supporting prescribed fire, managed wildfire, and Indigenous fire stewardship as targeted adaptation tools where ecological, cultural, and operational conditions are suitable^32,60,61^. Benefits ranked by area were greatest where low-severity burning is ecologically plausible and likely to reduce future severity without creating high-severity legacies, such as many Mediterranean and temperate conifer systems with increasing fire risk under a changing climate^1,48,55,62^. In contrast, many tropical moist broadleaf and boreal forests showed lower relative mitigation potential or greater risk of undesirable outcomes^63^, suggesting that low-severity fire management should not be generalized across all forest biomes. This biome variation is consistent with the stronger role of fire severity in structural recovery than in functional recovery (**Text S2** and **Fig. S2**): biomes where non-recovery is mainly structural tend to benefit more from severity mitigation driven by fuel removal, whereas function-limited systems may remain constrained by climate, water availability, or productivity even after severity is reduced (**Fig. S15**).

Several caveats qualify these conclusions. First, recovery in this study denotes the return of remotely sensed structural or functional indicators to contemporary pre-fire baselines; it does not confirm recovery of species composition, stand age structure, or ecosystem state, which can follow different trajectories ^6,16,64^. Second, MODIS NPP and ET estimates rely on plant-functional-type parameterizations that may not fully capture post-fire shifts in vegetation type or plant traits, adding uncertainty to functional recovery estimates^65,66^. Third, the approximately two-decade satellite record constrains inference for recent fires and slow-growing forests because recovery trajectories require long post-fire observation windows, and late-period MODIS observations may be affected by changes after orbit-maintenance^6,67,68^. Fourth, apparent non-recovery may also reflect fire severity interacting with a changing climate that no longer supports pre-fire biomass, canopy structure, or tree regeneration^69^. Finally, prior-fire mitigation estimates are statistical counterfactuals that should be interpreted as screening-level evidence that requires testing with additional global field observations and daily fire-weather data, as well as local fire-behavior constraints, and place-based management knowledge^11,70^.

Overall, our study advances a consequence-based view of wildfire risk: the most important fires are not necessarily those that burn the largest area, but those after which forests are least likely to recover to pre-fire status. By linking individual fire events to recovery trajectories, the crucial fire framework provides a global screening tool for identifying where wildfire is likely to cause persistent structural or functional loss ^3,22,71,72^ and where intervention may most effectively reduce long-term ecological consequences. For management, the main contribution is to identify where prescribed fire, managed wildfire, Indigenous fire stewardship, or other low-severity fire practices are most likely to reduce future severity and recovery-threshold risk, rather than treating fire use or fire exclusion as universal strategies^7,8^. Future work integrating field monitoring, fire-behavioring modeling, local fire knowledge, and higher-resolution recovery data can translate these global priorities into operational guidance for specific landscapes.

## Methods

### Dataset and pre-processing

We integrated the following datasets in a four-step workflow organized around individual fire events as the primary unit of analysis (**Extended Data Fig. 7**). First, during pre-processing, FIRED ^73^ fire perimeters were combined with MODIS land cover ^74^ and the Copernicus 100-m cropland layer ^75^ to retain forest fire events and reduce contamination from direct land-use conversion. Second, annual MODIS vegetation products ^76–79^ were used to reconstruct post-fire recovery trajectories in forest structure and ecosystem function, from which we identified persistent forest loss. Third, event-level recovery outcomes were linked to fire attributes, pre-fire vegetation condition, climate ^80^, and topography ^81^ to classify crucial fires with high probability of persistent non-recovery. Finally, prior-fire history was used to estimate the potential mitigation effect of low-severity fire, and the GRIP4 road network ^82^ was used only at the prioritization stage to identify road-accessible landscapes where management interventions may be operationally feasible.

#### Fire event dataset and fire severity definition

To identify individual wildfire events and their attributes at the global scale, we used the Fire Event Delineation (FIRED) dataset ^35,73^. FIRED is derived from the MODIS Burned Area product (MCD64A1), which provides monthly burn dates at 500 m resolution. By aggregating burned pixels that are contiguous in space and time, FIRED converts pixel-level burn observations into discrete fire-event perimeters. We used the global archive from 2001 to 2024 and retained events occurring primarily in forested landscapes. For each event, we extracted the start date, end date, and final burned area, yielding a global geodatabase of approximately 3,087,000 forest fire events for analysis. Fire severity was quantified as the proportional reduction in vegetation condition from the pre-fire state to the post-fire state:

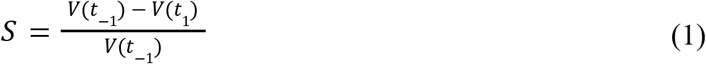

where *S* is fire severity, *V*(*t* _-1_) is the vegetation indicator value (TC, LAI, NPP, or ET) in one year before fire, representing the pre-fire baseline, and *V*(*t* _1_) is the value in the first post-fire year, representing the maximum observed fire impact. This normalization expresses severity as the fraction of pre-fire vegetation condition lost immediately after fire, with larger values indicating greater fire-induced decline and values near zero indicating little change.

#### Forest mask

To define forest extent and separate forest fires from non-forest fires, we used the annual MODIS Land Cover Type product (MCD12Q1) ^74^. Forest pixels were identified using the IGBP forest classes (classes 1 to 5) in the year preceding each fire. An event was retained as a forest fire when at least 30% of its burned pixels were classified as forest. Forest burned area and recovery trajectories were then calculated only for these pre-fire forest pixels, rather than for the entire event perimeter. MODIS annual land cover was not used to infer post-fire land-cover transitions. For biome-scale and ecoregion-scale analyses of crucial fire trends, we aggregated results using the WWF Terrestrial Ecoregions of the World classification^41^. Because WWF biomes represent broad ecological regions rather than pixel-level land cover, biomes such as tropical grasslands and savannas can contain forest patches; estimates reported for these biomes include only MODIS-defined pre-fire forest pixels.

#### Land-use change fire exclusion

To minimize contamination from known land-use fires and conversions, we applied independent land-use datasets. In the Brazilian Amazon, we used MapBiomas Brazil to exclude events in which more than 50% of valid burned pixels were mapped as agricultural or other excluded land uses^83^. Globally, we supplemented this filter with GLAD cropland maps for 2003, 2007, 2011, 2015, and 2019^84^. For each fire year, we used the pixel-wise maximum of the first three maps available in or after that year, with the 2019 map used for fires after 2019, and excluded events when more than 50% of valid pixels were mapped as cropland. Events meeting either regional or global exclusion criterion were removed. Because these products may not fully capture pasture expansion, shifting cultivation, or plantation establishment, the filters minimize rather than eliminate contamination from land-use conversion. Thus, persistent forest loss refers to non-recovery of pre-fire forest structure or function after filtering known land-use conversions, rather than to direct conversion itself. The land-use filters excluded 7.5% of the globally mapped persistent-loss area, including most mapped losses in the Amazon and smaller portions in Africa and Southeast Asia (**Fig. S17**)

#### Structure and function indicators for forest

To quantify the divergence between structural and functional recovery, we acquired a suite of MODIS-derived land products^76–79^. All vegetation data were processed at (or resampled to) a 500-m spatial resolution to match the fire event perimeters. (1) To quantify post-fire recovery of forest structure, we used two MODIS-derived indicators at 500 m resolution. Annual Percent Tree Cover was obtained from the Vegetation Continuous Fields product (MOD44B) as a measure of canopy closure, and represents the percentage of each pixel covered by tree canopy, ranging from 0 to 100%. Leaf Area Index (LAI) was obtained from the 8-day composite product MOD15A2H. To reduce seasonal noise and residual cloud effects, we summarized annual LAI using the 95th percentile of 8-day observations within each year, representing peak canopy condition. (2) To quantify post-fire recovery of ecosystem function, we used two complementary MODIS-derived indicators related to carbon uptake and water flux. Annual Net Primary Productivity (NPP) was obtained from MOD17A3HGF as a measure of ecosystem carbon accumulation. Annual Evapotranspiration (ET) was obtained from MOD16A3GF to represent ecosystem water use. Together, these four indicators allowed us to compare post-fire recovery in forest structure and ecosystem function within a consistent satellite-based framework.

#### Environmental and topographic drivers

To represent environmental conditions that may influence post-fire recovery beyond fire characteristics, we integrated climate and topographic data. Monthly climate variables were obtained from TerraClimate^80^, which provides gridded climate and water-balance data at approximately 4-km spatial resolution, and the resulting climate metrics were bilinearly resampled to the 500-m MODIS grid. For each fire event, we first aggregated monthly TerraClimate variables to annual values for each of the first five post-fire years, then averaged those five annual values to characterize early post-fire climate. The variables included annual precipitation, annual maximum temperature, annual mean Palmer Drought Severity Index (PDSI), and annual mean soil moisture, which were resampled to the 500-m MODIS grid using bilinear interpolation. Topographic variables were derived from GMTED2010^81^. Elevation was used to account for terrain-related constraints on regeneration under the 500-m resolution grid.

#### Road-based accessibility and mitigation-priority ranking

To approximate where fire-management interventions may be operationally accessible, we used the Global Roads Inventory Project version 4 (GRIP4) vector road dataset^82^. We reprojected GRIP4 roads to the 50-km equal-area hexagon grid and classified a hexagon as accessible when at least one road segment intersected it. Accessibility was used to constrain the mitigation-priority ranking: inaccessible hexagons were retained in the map but excluded from ranking. Among road accessible hexagons with a nonzero realized mitigation area, we ranked hexagons in descending order of mitigation area. Percentile classes were based on the number of ranked hexagons, with the top 1% representing the highest-ranked 1%, the top 2–10% representing the next 9%, and the remaining 90% representing all lower-ranked hexagons.

### Recovery potential identification

To enable a standardized comparison of post-fire trajectories across disparate global biomes and baseline productivity levels, we conducted a time-series analysis based on relative recovery proportion for each fire event. For each burned forest pixel within a fire perimeter, we quantified annual recovery status at time *t_n_*, where *n* denotes years after fire, as the fraction of the initial fire-induced loss that had been regained relative to the pre-fire baseline:

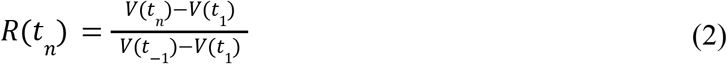

where *R*(*t* _n_) is the recovery proportion at year *t_n_* post-fire, *V*(*t* _-1_) is the value in the first post-fire year, representing the maximum observed fire impact, and *V*(*t*_-1_) is the pre-fire baseline, calculated from the year before fire. This normalization places all recovery trajectories on a common scale, where 0 represents the immediate post-fire state, 1 indicates return to the pre-fire baseline, and values above 1 indicate recovery beyond the pre-fire level. By tracking this normalized value annually, the analysis captures the temporal pathway of post-fire recovery of vegetation conditions (**Fig. 1A&B** and **Fig. S1**).

Post-fire recovery was modeled as a successional regrowth process following disturbance. In disturbance ecology, burned areas can be viewed as dynamic patches in which vegetation loss is followed by establishment, regrowth, community turnover, and potential reorganization, requiring recovery to be analyzed as a time series rather than a single post-fire condition^85^. Forest recovery models also recognize that regrowth is constrained by an upper recovery potential, with biomass or canopy attributes increasing after disturbance and eventually approaching a saturation level ^71^. By comparing seven growth models (**Text S1**), we therefore fitted a Gompertz growth function^86^ to the observed event-level *R*(*t* _n_) values as a function of years after fire (*t_1_* to *t_n_* ), separately for each fire event and vegetation indicator. Depending on the fire year, each trajectory contained 9-23 annual post-fire observations through 2024:

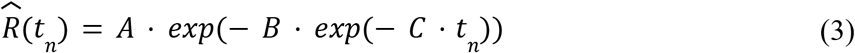

where *Y*(*t* _n_) represents the modeled recovery stage at year *t* _n_, *B* controls the displacement of the curve along the time axis, and *C* represents the intrinsic recovery rate. The upper asymptote *A* is used as the estimate of long-term recovery potential.

We used the fitted asymptote *A* to diagnose whether post-fire recovery was projected to return to the pre-fire baseline. Because the recovery time series was normalized to the pre-fire condition, 1 represents the pre-fire state, *A* ≥ 0. 95 was treated as near-complete recovery, allowing a 5% tolerance for remote sensing data noise, interannual variability, and model fitting uncertainty.Conversely, *A* < 0. 95 indicates that the fitted trajectory stabilizes below the pre-fire baseline and interpreted as persistent non-recovery, as long-term degradation or transition toward an alternative vegetation state. We used 0.95 as the primary operational threshold because it avoids requiring exact return to baseline while still representing a standard for recovery. We tested the robustness of this choice by repeating the analysis across recovery thresholds from 0.80 to 1.00, and threshold selection does not impact spatial pattern of persistent loss (**Text S3** and **Fig. S10**).

NPP and ET vary strongly from year to year because of regional climate fluctuations, even in forests that have not burned ^87^. To reduce this background variability, we normalized NPP and ET using nearby unburned forests as annual references. For each region, year, and indicator, we identified unburned forest pixels as MODIS forest pixels with no recorded fire in the annual fire data. These pixels were aggregated to 1° × 1° grid cells after reprojection to geographic coordinates. Within each grid cell, we calculated the mean NPP or ET of unburned forests for each year and compared it with the long-term mean across all years. The ratio between the long-term mean and the annual mean was used as a correction factor for that grid cell and year. We then multiplied burned-pixel NPP and ET values by this factor before estimating recovery trajectories. This correction removes broad annual climate-driven fluctuations while preserving local post-fire recovery dynamics. This normalization was applied only to NPP and ET, not to tree cover or LAI.

### Crucial fire identification

Using the recovery outcomes derived from modeled recovery potential as binary response labels, we trained indicator-specific XGBoost classifiers to estimate the probability of post-fire recovery for tree cover, LAI, NPP, and ET ^88^. Each record represented one fire event evaluated for one vegetation indicator. Predictors included relative fire severity, fire size (after log-transformed), pre-fire indicator value, elevation, maximum temperature, precipitation, PDSI, and soil moisture, after multicollinearity check. After excluding invalid recovery fits, negative or very low severity events (<0.1) below the analysis threshold, and records with missing predictors, the dataset contained approximately 590,000 event-indicator records across training and validation data.

For each indicator, records were split into stratified training and validation sets, with 70% used for training and 30% retained for validation. Hyperparameters were optimized using randomized search with three-fold stratified cross-validation. We report validation AUC as the primary measure of model skill because it evaluates discrimination across probability thresholds. AUC ranged from 0.75 to 0.83 across indicators: TC = 0.76, LAI = 0.75, NPP = 0.83, and ET = 0.79; validation accuracy ranged from 0.69 to 0.76. For each indicator, a fire was classified as a crucial fire when its predicted recovery probability was <0.25, equivalent to a modeled non-recovery probability >0.75. An event was classified as an overall crucial fire if it met this criterion for at least one structural or functional vegetation indicator. The sensitivity analysis for crucial fire threshold (**Text S3**) shows there is no sudden change with threshold change, and spatial pattern remains stable.

We interpreted the fitted models using gain-based feature importance and partial dependence analysis. To estimate uncertainty, the best model for each indicator was refitted 100 times with different random seeds, and feature importance and partial dependence curves were summarized across runs. We focused on fire attributes because they provide the clearest management leverage: climate and topography are largely constant within a landscape, whereas fire severity and, to a lesser extent, fire size can be altered before or during fire through fuel management, prescribed fire, or suppression. Across all vegetation indicators, fire severity was consistently among the strongest predictors (**Fig. 2C** and **Fig. S2**), and recovery probability declined sharply with increasing severity. Fire size had a weaker and more gradual effect.

For spatial and temporal analysis, crucial fire predictions were aggregated to a 50-km equal-area hexagon grid. Forest burned area for each event was calculated from the forest fraction within its perimeter, and annual crucial fire area was summed within each hexagon. To reduce fine-scale spatial variability, annual crucial-fire area was smoothed using Gaussian distance weights across neighboring hexagons within four center-to-center grid spacings, with σ set to half the neighborhood radius. We estimated temporal trends in the annual crucial fire area using Theil–Sen slopes, with Kendall’s rank correlation used to assess significance. For global maps and biome summaries, the overall crucial fire area was defined using the union of indicator-specific predictions, so that a fire was counted as crucial if at least one structural or functional recovery model classified it as high risk.

### Counterfactual analysis of prior fire mitigation

Because fire severity was a major predictor of crucial fire occurrence (**Fig. S2**), prior low-severity fire represents a potential management pathway. Prior low-severity fire may reduce crucial fire risk by lowering fuel continuity and modifying stand conditions, thereby increasing local resistance to severe reburning. We tested whether prior low-severity fire reduced the severity of subsequent wildfires and the modeled risk of persistent forest loss. Because prior fire and reburn (subsequent fire) do not completely coincide, we used an observational counterfactual design to compare burned locations with and without prior-fire history under similar conditions. Pixels that burned in a focal wildfire and had also burned 2–10 years earlier were treated as prior-fire pixels ^9^. For each vegetation indicator and biome, prior low-severity fire was defined as a previous burn with severity at or below the 25th percentile of the corresponding severity distribution (**Fig. S5**); pixels with higher prior-fire severity were retained as a comparison group.

For each prior-fire pixel, we selected a control pixel that burned in the same focal wildfire but had no recorded prior fire in the 2–10 year window. Matching was constrained within the same fire event and biome and was based on standardized pre-fire indicator value and spatial proximity within the fire footprint. This design compared locations exposed to the same subsequent fire while reducing differences in pre-fire vegetation condition and local setting. An example matched pair is shown in **Extended Data Fig. 2**, where a prior-burn pixel and a nearby no-prior-burn control are identified within the same wildfire, together with their pre-fire and post-fire LAI conditions.

We then estimated how prior-fire history modified subsequent fire severity using generalized additive models. For each vegetation indicator and biome, we fitted separate severity curves for control pixels, pixels with prior low-severity fire, and pixels with higher-severity prior fire as a function of pre-fire vegetation condition. The mitigation effect of prior low-severity fire was defined as the difference between the fitted control and prior low-severity curves:

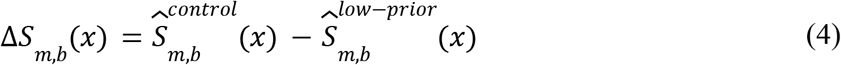

where *x* is the pre-fire indicator value, *m* is the vegetation indicator, and *b* is the biome. Positive values Δ*S* indicate reduced subsequent fire severity relative to matched controls; negative values indicate increased severity. The higher-severity prior-fire group was used to test whether this effect was specific to low-severity prior fire. The fitted severity relationships and their differences are illustrated in **Extended Data Fig. 4**: panel A shows the LAI-based hexagon-density plots and GAM curves for the three prior-fire-history groups, panel B shows the estimated low-severity prior-fire effect across TC, LAI, NPP, and ET, and panel C shows the corresponding comparison for higher-severity prior fire.

Finally, we translated the estimated severity reduction into changes in crucial fire risk using the XGBoost recovery models described above. For each event-indicator record, we created a low-severity prior-fire scenario by reducing observed fire severity according to Δ*S _m,b_*(*x*), while holding fire size, pre-fire vegetation, climate, and topography constant. The adjusted severity was then passed through the corresponding XGBoost model to estimate counterfactual recovery probability. A fire was considered mitigated for a given indicator when it was classified as crucial under observed conditions but not under the low-severity prior-fire scenario. Mitigated forest burned area was then summed globally and aggregated to 50-km equal-area hexagons to map the spatial density of potential crucial fire reduction. Furthermore, road data was included to detect accessible hexagons; it did not influence recovery estimation or crucial fire classification, but provided a practical accessibility constraint for mapping management opportunities.

## Data availability

FIRED fire-event perimeters were generated using FIREDpy from the MODIS MCD64A1 burned-area product (https://github.com/earthlab/firedpy). MODIS MCD64A1, MCD12Q1 (land cover map), MOD44B (tree cover), MOD15A2H (LAI), MOD17A3HGF (NPP) and MOD16A3GF (ET) data were obtained from the NASA Land Processes Distributed Active Archive Center (https://search.earthdata.nasa.gov/search/). Cropland data were obtained from the GLAD global cropland dataset (https://glad.umd.edu/dataset/croplands), and Brazilian land-cover data were obtained from MapBiomas Brazil, Collection 11 (https://plataforma.mapbiomas.org/projects/mapbiomas/brazil). Climate data were downloaded from TerraClimate (https://www.climatologylab.org/terraclimate.html), elevation data from GMTED2010 (https://earthexplorer.usgs.gov/), and road data from GRIP4 (https://doi.org/10.5281/zenodo.6420961). Raw FIA data were downloaded from the FIA DataMart (https://research.fs.usda.gov/products/dataandtools/fia-datamart).

## Code availability

Code and purpose-limited data are available in Figshare, and publicly available once the article is accepted.

## Acknowledgement

This work was supported by the Life-cycle Assessment Synthesized with Ecosystems and Risk (LASER) project. S.G., S.Z, Y.Z, and K.Z. received support from NSF (grant no. 2306198) and USDA McIntire-Stennis Capacity Grant (award no. 25-PAF01509). J.A.W received support from NASA Terrestrial Ecology, NASA Land Cover Land Use Change, and NASA Early Career Investigators Program. We thank Dr. Jennifer Balch for providing the FIRED dataset and answering methodological questions; Kylie Liang and Yuxin Yao for assistance with collecting and processing MODIS and fire data; and Dr. Dimitrios Gounaridis, Neal Harbaugh and members of the Zhu Lab for suggestions on figure presentation and comments on the manuscript.

## Author contribution

S.G., B.P.G., and K.Z. conceptualized this study. S.G. and K.Z. developed the conceptual framework for evaluating persistent post-fire forest loss and identifying crucial fires. S.G. and S.Z. developed the forest-recovery trajectory analysis, crucial-fire classification models, and associated software. S.G. designed and conducted the prior-fire matching analysis and model-based counterfactual assessment of low-severity fire mitigation. J.T performed the FIA-based validation. S.G., S.Z. and Y.Z. produced the figures and visualizations. J.A.W., Z.Z., and T.G. contributed expertise in remote sensing, forest ecology, wildfire dynamics, and agriculture fire exclusion. M. O., Y. L., and P. B. R. contributed expertise in fire-mitigation assessment, climate-condition analysis, uncertainty quantification and sensitivity analysis. S.G., S.Z. and Y.Z. interpreted the results, and S.G., wrote the first draft of the manuscript. P. B. R., B.P.G. and K.Z. supervised the research, and B.P.G. and K.Z. acquired funding. All authors contributed to interpreting the results and reviewing and editing the manuscript.

---

**Extended Data Figure 1.**
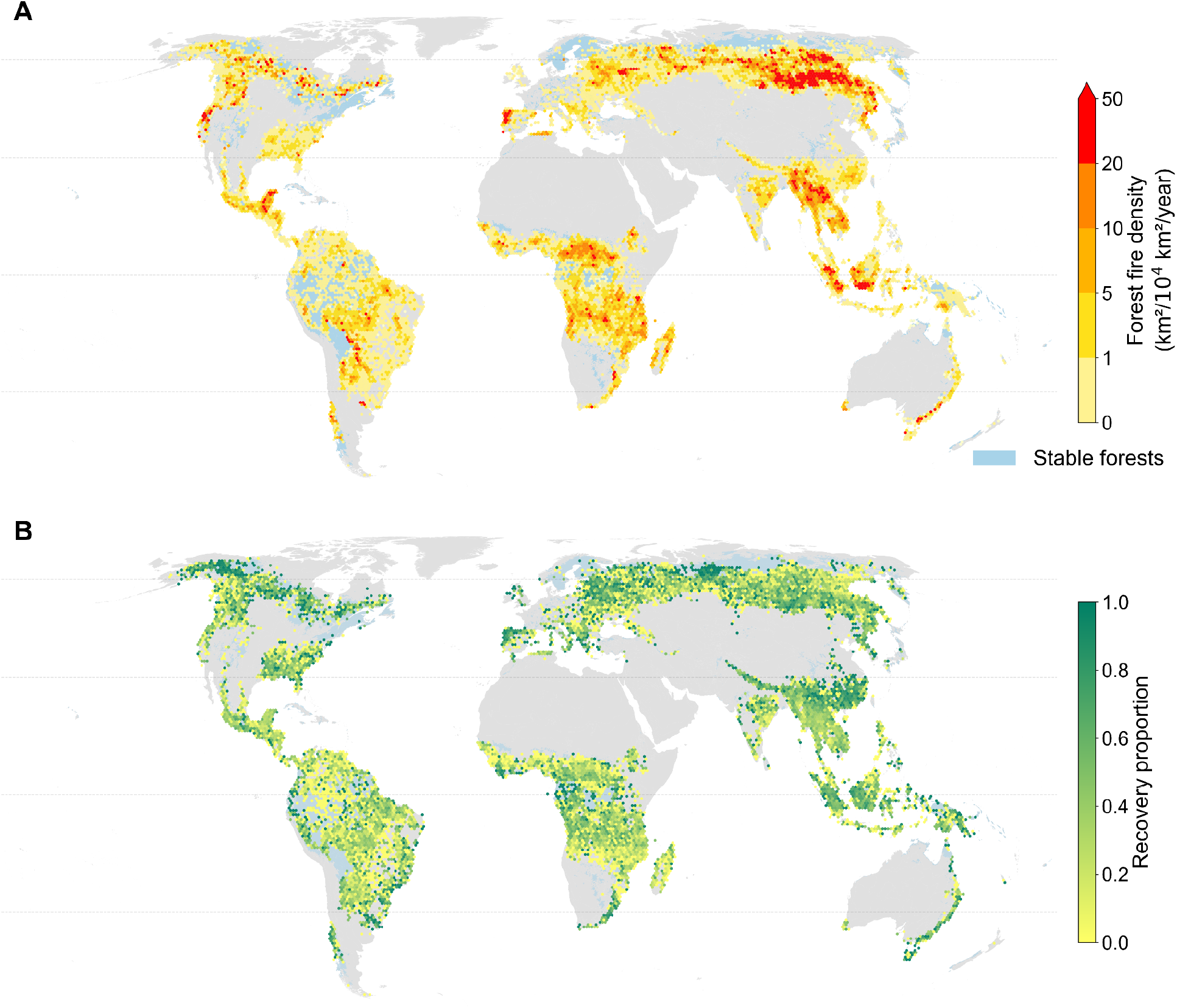
Global forest fire density and postfire recovery proportion. (A) Global forest fire density summarized on 50-km equal-area hexagons for fires occur during 2001–2014. Warm colors indicate higher annual fire-affected forest area, expressed as km² /10⁴ km² /year, whereas blue denotes stable forests without recorded fire. (B) Global projected recovery proportion of fire-affected forests, defined as the fraction of burned forest area within each hexagon that recovered toward prefire conditions. Higher values indicate dominance of recovered forests, whereas lower values indicate a larger share of persistent postfire forest loss.

**Extended Data Figure 2.**
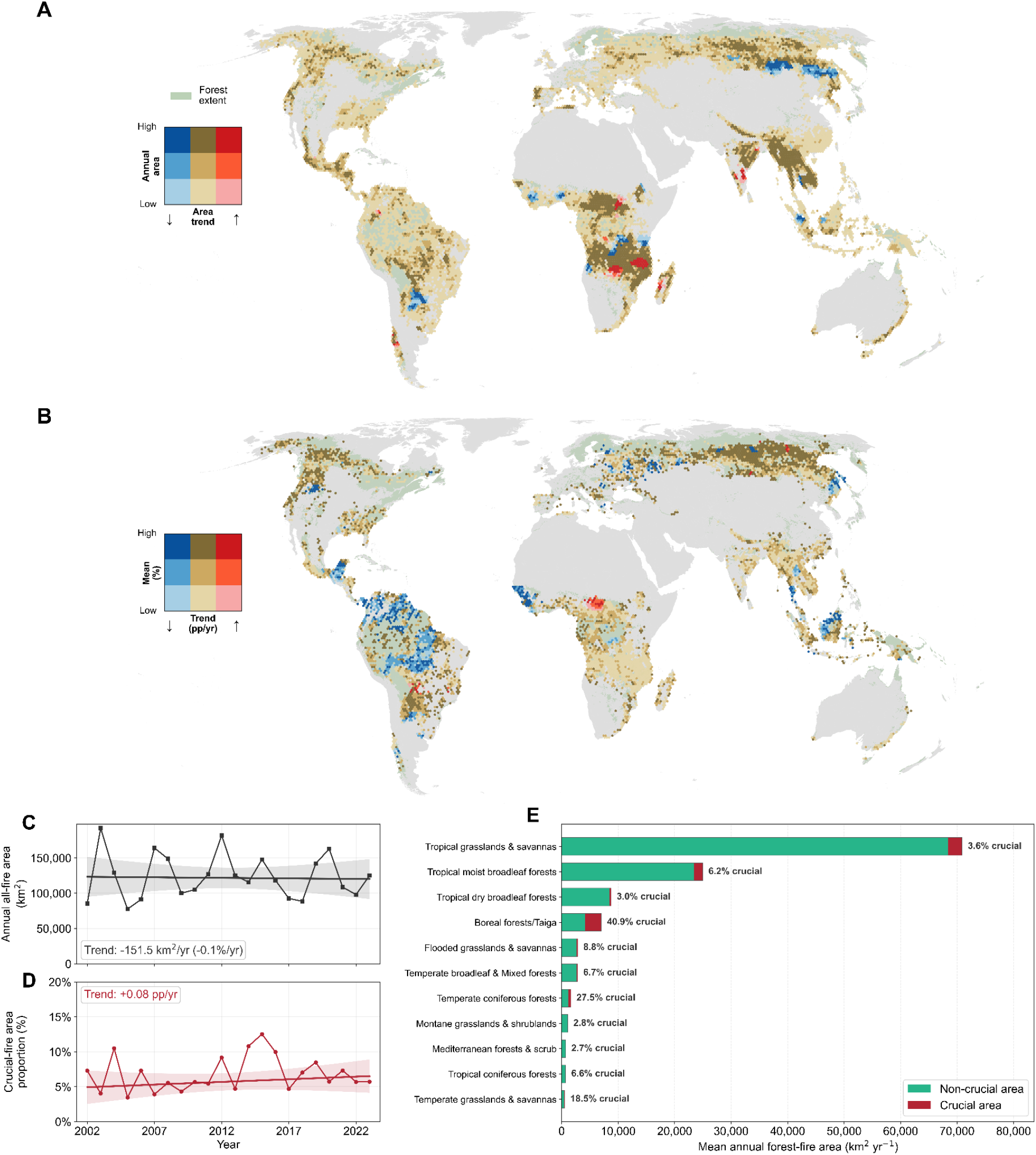
Global patterns of all and crucial forest fires, 2002–2023. Crucial fires were defined as events with a modeled recovery probability <0.25 for at least one ecosystem metric: TC, LAI, NPP, or ET. (A) Mean annual all-fire area density and its temporal trend across 50-km-side hexagons. (B) Mean crucial fire area proportion and its temporal trend; only hexagons containing both all-fire and crucial fire area are shown. In A and B, darker colors indicate greater magnitude, blue and red indicate decreasing and increasing trends, respectively, and tan/brown indicates weak or nonsignificant trends. Pale green denotes forest extent. Spatial trends were estimated from Gaussian-smoothed annual values using Theil–Sen slopes and Kendall tests. (C) Global annual all-fire area. (D) crucial fire area as a percentage of annual all-fire area. Points show annual values, fitted lines show Theil–Sen trends, and shading indicates approximate 95% uncertainty envelopes. (E) Mean annual forest-fire area by WWF biome, partitioned into non-crucial and crucial area; labels indicate the percentage classified as crucial.

**Extended Data Figure 3.**
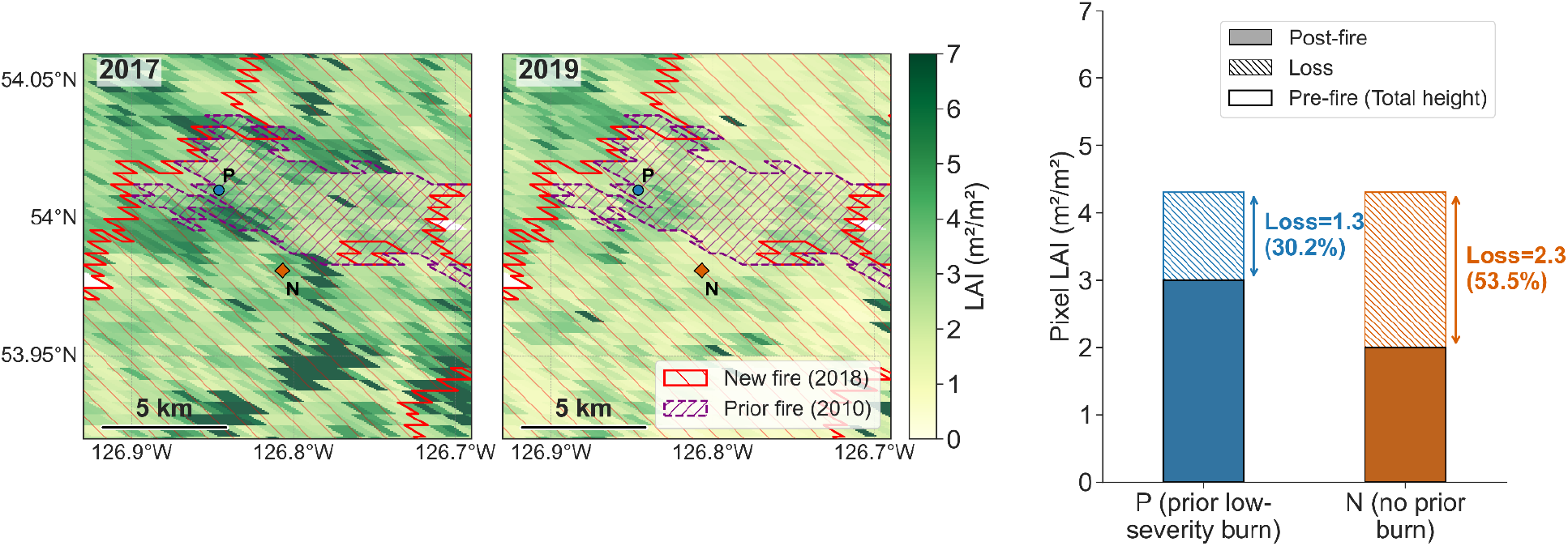
Example of paired-pixel comparison used in the prior-fire mitigation analysis. The current fire occurred in 2018 (red outline), while the prior fire occurred in 2010 and is shown as the purple dashed perimeter. Within a focal wildfire, a pixel with prior low-severity fire history (P) is matched to a nearby pixel without prior fire history (N). Pre-fire (2017) and post-fire (2019) LAI maps show that, despite similar initial canopy conditions, the prior-burn pixel experienced smaller LAI loss following the 2018 fire, illustrating how antecedent mild burning can reduce subsequent fire impact.

**Extended Data Figure 4.**
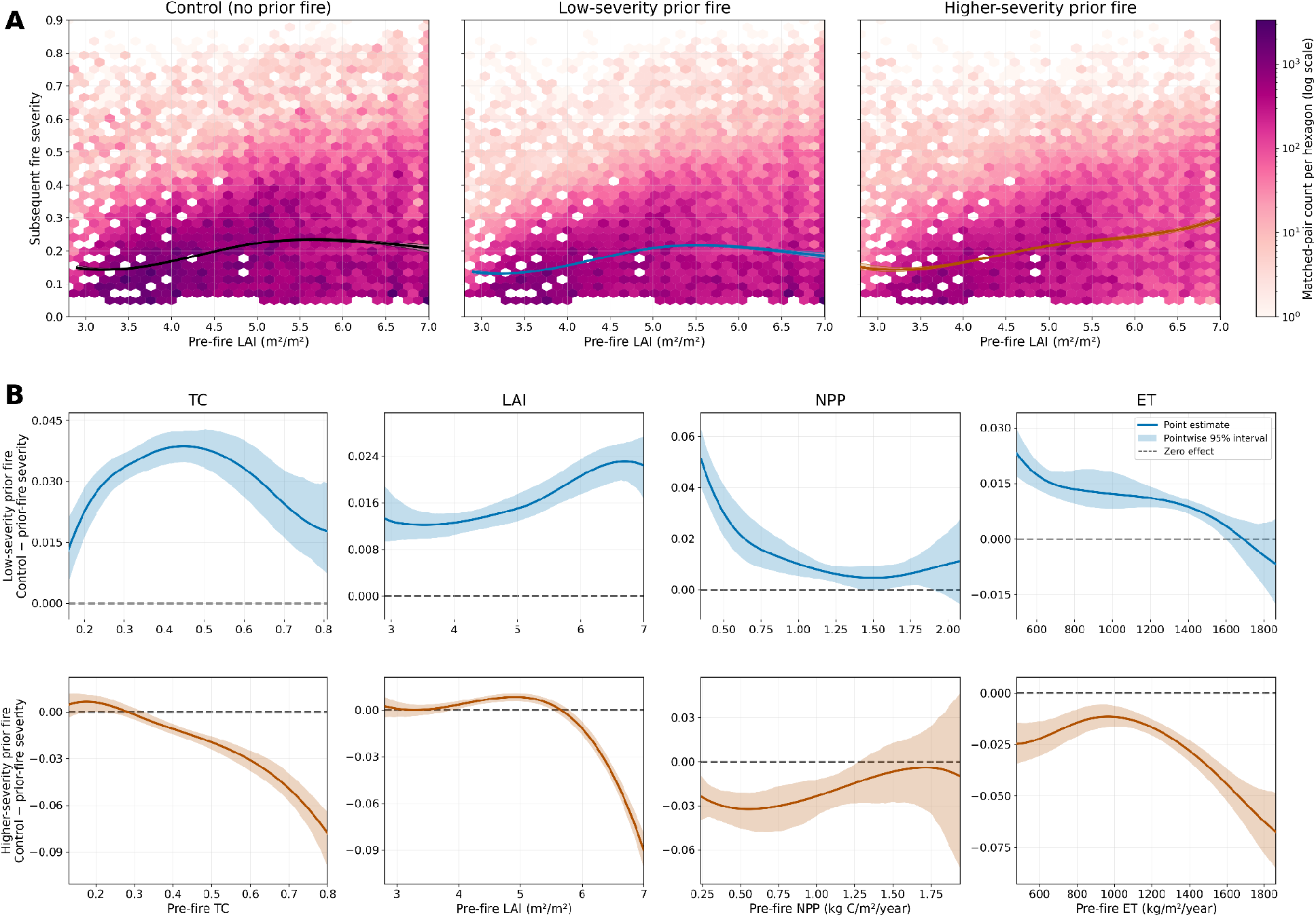
Prior-fire history modifies the relationship between subsequent fire severity and pre-fire vegetation condition, with uncertainty quantified by a fire-cluster bootstrap. (A) Hexagon-density plots of subsequent fire severity versus pre-fire leaf area index (LAI) for matched control pixels without prior fire, pixels with a low-severity prior fire, and pixels with a higher-severity prior fire, pooled across biomes. Hexagon colors indicate matched-observation density on a logarithmic scale. Black, blue, and brown curves show the respective generalized additive model estimates, and bands of the same color show pointwise 95% intervals from 1,000 fire-cluster-bootstrap replicates. Curves and densities are shown over their common uncertainty-supported LAI range. (B) Estimated prior-fire effects across tree cover (TC), LAI, net primary productivity (NPP), and evapotranspiration (ET). The upper row shows low-severity prior fire and the lower row shows higher-severity prior fire. Effects are expressed as control minus prior-fire subsequent severity; positive values indicate mitigation, whereas negative values indicate exacerbation. Solid curves show point estimates, matching-color bands show pointwise 95% fire-cluster-bootstrap intervals, and dashed horizontal lines mark zero effect..

**Extended Data Figure 5.**
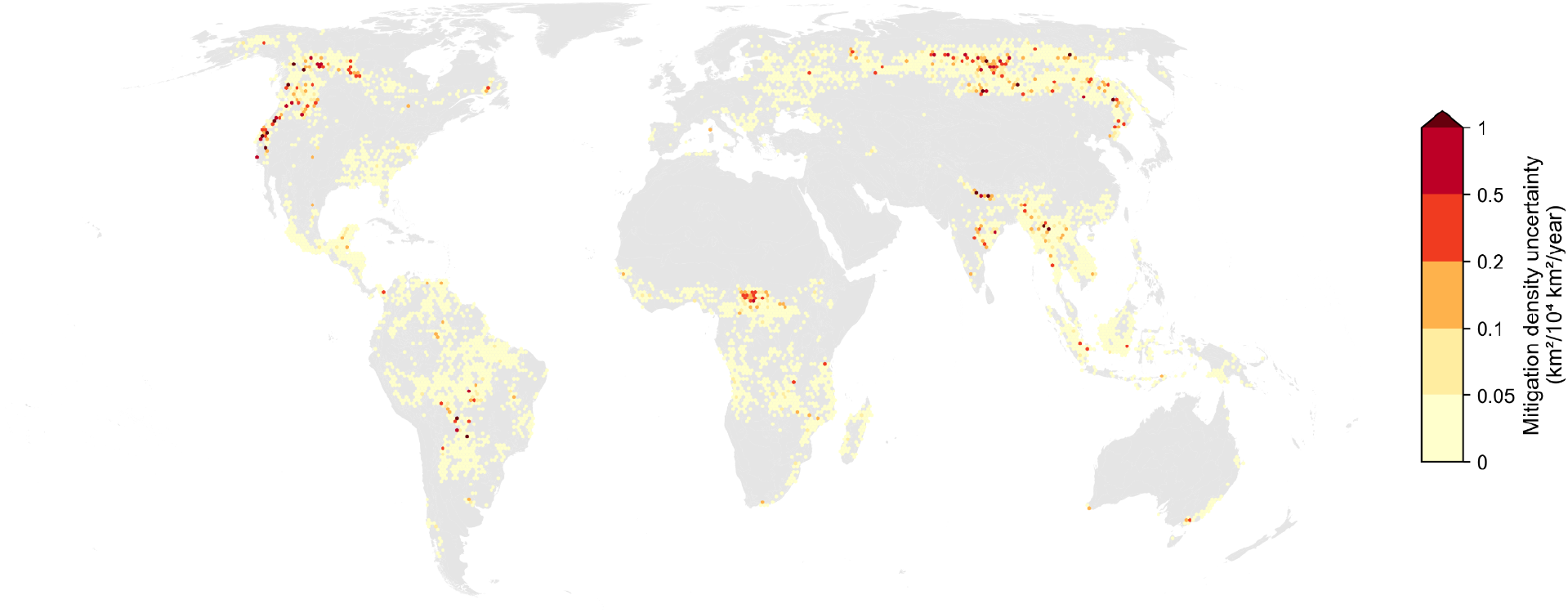
Spatial uncertainty in estimated prior-fire mitigation density. Absolute uncertainty in the mitigation density associated with prior low-severity fire, summarized on 50-km equal-area hexagons. For each of 1,000 bootstrap realizations, biome-specific mitigation curves were propagated through the fixed XGBoost recovery models. Fires with an observed recovery probability below 0.25 were classified as crucial, and those reaching a counterfactual probability of 0.25 or higher were classified as mitigated. Mitigated forest-fire area was averaged across TC, LAI, NPP, and ET and normalized by the 23-year observation period. Uncertainty was quantified as the standard deviation of annual mitigation density across curve realizations and is expressed as km²/10⁴ km²/year; darker colors indicate greater uncertainty. Fire events, locations, and years were held fixed, so the map represents uncertainty arising from the estimated mitigation curves rather than interannual fire variability or recovery-model parameter uncertainty.

**Extended Data Figure 6.**
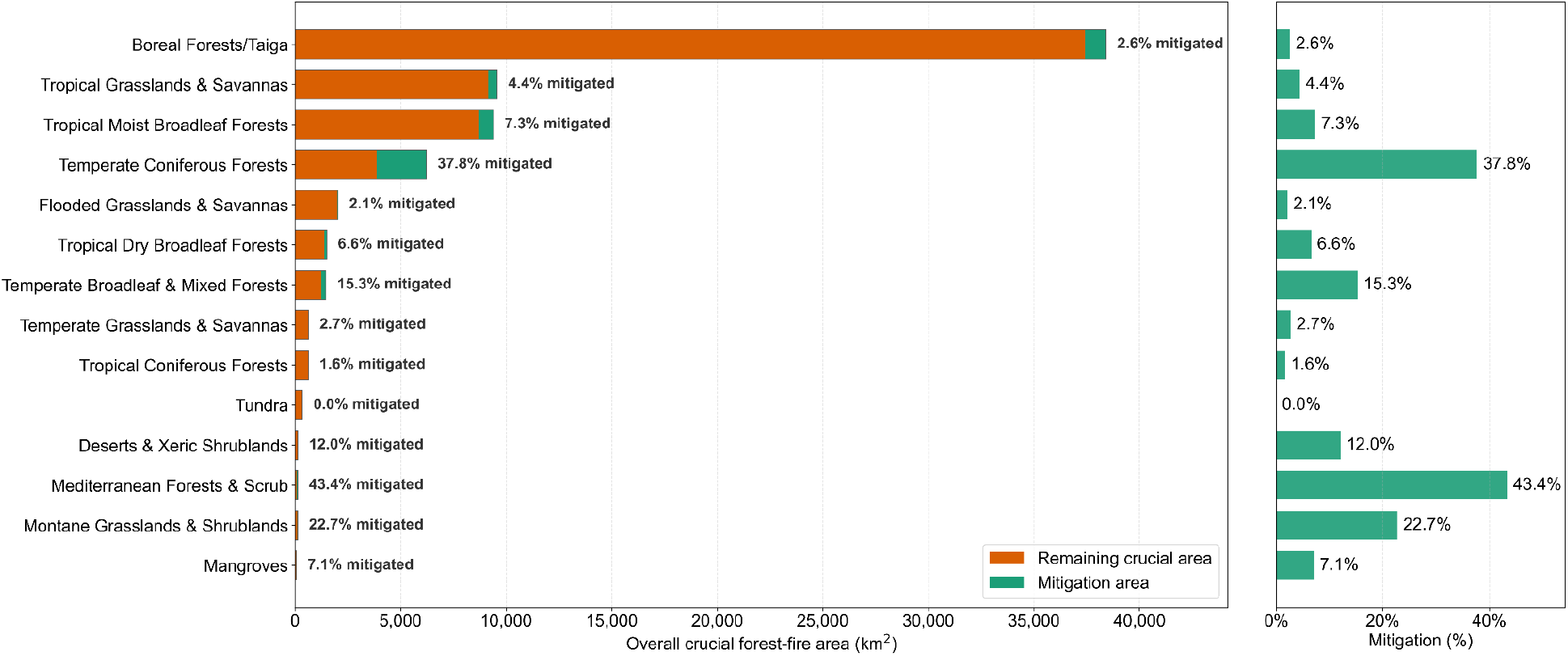
Biome-level mitigation potential under a low-severity prior-burn scenario. Biome-level estimates of absolute and relative mitigation potential for crucial forest-fire area. The left panel partitions overall crucial forest-fire area into remaining crucial area and mitigation area under the low-severity prior-burn scenario. The right panel shows the corresponding mitigation percentage for each biome, calculated as the proportion of overall crucial forest-fire area that could be mitigated.

**Extended Data Figure 7.**
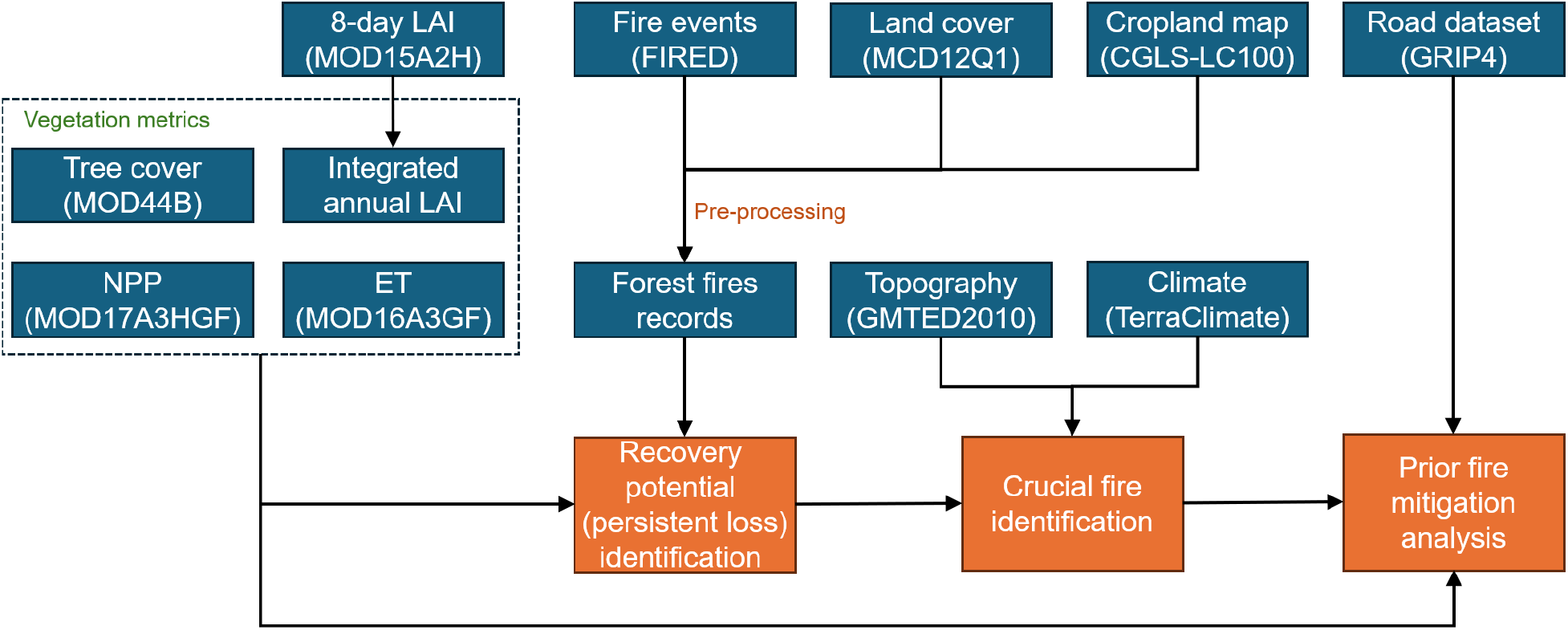
Dataset integration and processing workflow for global forest-fire recovery analysis. Fire-event, land-cover, vegetation, climate, topographic, cropland, and road-network datasets were integrated to support four analytical steps: forest-fire event screening, post-fire recovery and persistent-loss identification, crucial fire classification, and prior-fire mitigation prioritization. Road data were used only to constrain the final management-priority analysis to accessible landscapes.

**Extended Data Figure 8.**
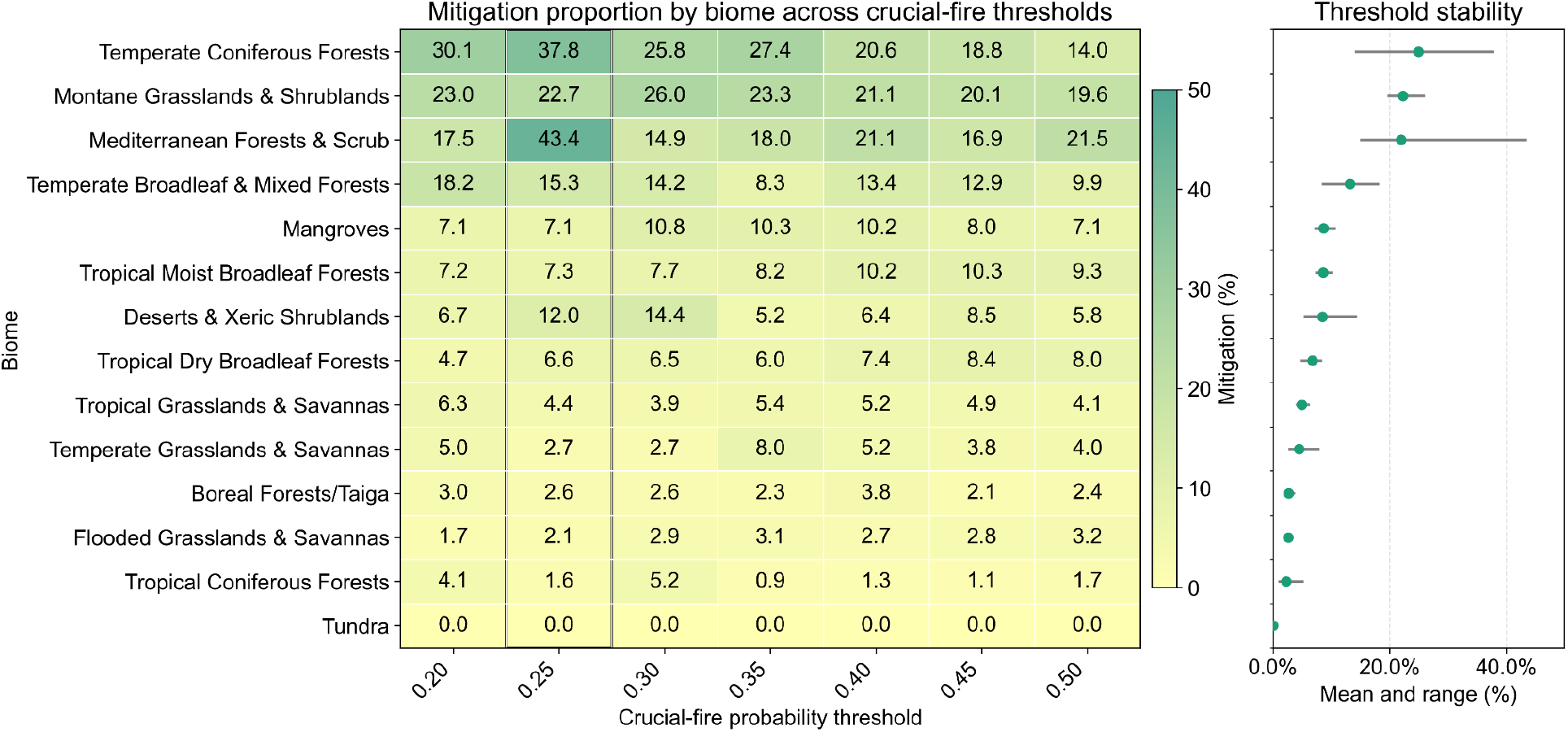
Sensitivity of estimated mitigation effects with different crucial fire probability thresholds. Biome-level mitigation proportions were recalculated across crucial fire probability thresholds from 0.20 to 0.50. The heatmap shows mitigation proportion for each biome and threshold, with darker colors indicating higher mitigation potential. The right panel summarizes threshold stability as the mean and range of mitigation proportion across thresholds for each biome.

