## Supplementary Information for "Prior low-severity fires reduce the risk of persistent forest loss from subsequent fires"

#### Text S1. Forest growth models selection

To estimate long-term post-fire recovery potential, we compared seven candidate growth models that represent common forms of ecological recovery trajectories, including linear recovery, saturating recovery, sigmoidal recovery, and power-law recovery[1–4]. All models were fitted to the normalized recovery proportion $Y(t_n)$, where $t_n$ represents n-year since fire and $Y(t_n)$=0.95 indicates return to the pre-fire baseline. The candidate models were:

| | | |
|---|---|---|
| Linear | $Y(t_n) = \alpha + \beta t_n$ | (1) |
| Logistic | $Y(t_n) = \frac{A}{1+e^{-k(t_n - t_0)}}$ | (2) |
| Monod | $Y(t_n) = A\frac{t_n}{h+t_n}$ | (3) |
| Chapman-Richards | $Y(t_n) = A(1 - e^{-kt_n})^m$ | (4) |
| Negative Exponential | $Y(t_n) = A(1 - e^{-kt_n})$ | (5) |
| Power | $Y(t_n) = ct_n^{\ b}$ | (6) |
| Gompertz | $Y(t_n) = A \cdot exp(-\ B \cdot exp(-\ C \cdot t_n))$ or $Y(t_n) = A \cdot exp(-\ exp(-\ k(t_n - t_0))$ | (7) |

where $A$ denoting the asymptotic recovery level for bounded models, $k$ is the recovery rate, $t_0$ is the inflection or lag-position parameter, $h$ is the half-saturation time, and $m$ as a shape parameter.

For each vegetation indicator: tree cover (TC), leaf area index (LAI), net primary productivity (NPP), and evapotranspiration (ET), we fitted all candidate models to the recovery time series and evaluated model performance using R2 and root mean square error (RMSE). Model performance was summarized across fire events. Across the four vegetation indicators, the Gompertz model provided consistently strong performance (**Table. S1** and **Fig. S9**), with mean $R^2$ of TC = 0.604, LAI = 0.648, NPP = 0.679, ET = 0.662 and mean RMSE of TC = 0.337, LAI = 0.284, NPP = 0.172, ET = 0.204. Its performance was comparable to or better than the other candidate models, particularly for recovery trajectories with an initial lag followed by rapid regrowth and eventual saturation.

We therefore used the Gompertz function for the main analysis because it combines good empirical fit with an ecologically interpretable asymptote. The fitted asymptote represents the projected long-term recovery potential relative to the pre-fire baseline, allowing each fire event to be classified according to whether recovery was projected to return to pre-fire conditions or stabilize below them. This choice was also conservative for the purpose of identifying persistent forest loss, because the model allows delayed recovery rather than assuming a linear or fixed-rate return to the pre-fire condition.

#### Text S2. Interpretation of structural and functional recovery

Our indicators represent complementary but imperfect axes of recovery. Tree cover is the most direct satellite proxy for whether tree canopy has returned, whereas LAI measures leaf area and NPP and ET measure ecosystem function[5]. Shrub or early-successional regrowth may partly or even strongly restore LAI, NPP or ET, but it should not generally restore the pre-fire tree-cover condition. Therefore, recovery in functional indicators without tree-cover recovery should be interpreted as functional green-up or early-successional recovery, not as confirmed mature-forest recovery [6]. Conversely, apparent tree-cover or LAI recovery does not guarantee recovery of the original species composition, biomass, age structure, biodiversity, or vertical complexity.

The contrast between structural and functional recovery helps explain why post-fire forest resilience cannot be inferred from greenness or canopy closure alone. In high-latitude forests, functional recovery often exceeded structural recovery, suggesting that early successional vegetation can restore part of the productivity and water-flux signal before mature forest structure returns. This pattern is consistent with evidence that post-fire tree regeneration can be constrained by large high-severity patches, limited seed availability, fire legacies, and post-fire drought, even where vegetation regrowth resumes[7–9]. In tropical and subtropical forests, tree cover or LAI more often approached pre-fire levels while NPP and ET remained depressed. This apparent structural recovery should be interpreted cautiously. Dense tropical canopies can saturate optical and LAI-based satellite signals[10,11], and recovered canopy cover may reflect pioneer or secondary vegetation rather than restoration of old-growth vertical complexity, biomass, species composition, or hydraulic function[12–15]. Thus, tropical forests may appear structurally recovered in coarse satellite indicators while remaining functionally or compositionally degraded.

#### Text S3. Sensitivity analysis

We tested whether the main conclusions depended on operational thresholds used to define persistent forest loss, crucial fires, and prior low-severity fire. These analyses were designed to evaluate the stability of spatial patterns and effect directions, rather than to identify an optimal threshold.

First, we varied the recovery threshold used to define persistent forest loss. The main analysis used ($A < 0.95$), where ($A$) is the fitted Gompertz asymptote and 1 represents full return to the pre-fire baseline. This threshold was chosen as a near-complete recovery criterion that allows a small tolerance around the baseline while avoiding the more permissive assumption that a 10% deficit still represents recovery. We repeated the calculation across thresholds from 0.80 to 1.00 at 0.05 intervals. As expected, stricter thresholds increased the area classified as persistent loss (from approximately 9,500 to 22,500 $km^2$/yr), but the broad spatial structure remained stable: major persistent-loss regions in western North America, the Amazon, central Africa, mainland Southeast Asia, and Siberia were retained across threshold choices (**Fig. S10**). The latitudinal contrast between structural and functional recovery also remained evident across thresholds, indicating that the main spatial and dimensional patterns were not artifacts of the default 0.95 cutoff.

Second, we tested the recovery-probability threshold used to classify crucial fires. The default classification defined a crucial fire as an event with predicted recovery probability $p<0.25$. We repeated the classification using two alternative thresholds that bracketed the default value: 0.20, 0.30, and 0.40 representing more conservative and more inclusive definitions of crucial fires, respectively. Although higher thresholds classified more burned forest area as crucial, the broad spatial and biome-level patterns were stable across thresholds (**Fig. S11**). In all three cases, crucial fire area remained regionally heterogeneous, with decreasing patterns in parts of the southeastern Amazon and increasing patterns in African savanna and savanna–forest regions. Biome-level summaries also retained the same redistribution pattern, with tropical moist broadleaf forests contributing much of the decrease and tropical grasslands and savannas contributing the largest increase. We further evaluated whether the same threshold choices altered the inferred suitability of biomes for prior-fire mitigation. Across thresholds, the highest mitigation proportions remained concentrated in a similar set of biomes, especially temperate coniferous forests, Mediterranean forests and scrub, and montane grasslands and shrublands, whereas boreal forests and tundra showed consistently low mitigation proportions (**Extended Data Fig. 8**). Thus, changing the crucial fire probability threshold affected the magnitude of estimated crucial fire area and mitigation, but did not substantially change the broad spatial redistribution pattern or the main biomes identified as high-benefit contexts for low-severity prior-fire management.

Third, we evaluated the threshold used to define prior low-severity fire. In the main analysis, prior low-severity fire was defined as prior fire severity at or below the 25th percentile. We varied this threshold from 0.10 to 0.50 at 0.05 intervals and refitted the severity-response curves. Across all tested thresholds, the mean difference between matched controls and prior low-severity fire remained positive for TC, LAI, NPP, and ET, indicating that prior low-severity fire consistently reduced subsequent fire severity (**Fig. S12**). The magnitude of the reduction declined as the threshold increased, which is expected because more moderate-severity prior burns were progressively included in the low-severity group. In contrast, higher-severity prior fire did not show a stable protective effect and often produced equal or higher subsequent severity than matched controls (**Fig. S13**). These results support the interpretation that mitigation is specific to low-severity fire legacies rather than prior burning in general.

Finally, we tested the sensitivity of prior-fire mitigation to the time window used to define prior burns. The main analysis used prior fires occurring 2 to 10 years before the focal fire. We fixed the minimum interval at 2 years and varied the maximum interval from 5 to 15 years. The estimated mitigation effect remained positive across all indicators and all tested windows (**Fig. S14**). TC showed a nearly constant severity reduction across windows, NPP showed a modest decline with longer windows, and LAI and ET showed stronger reductions when older prior fires were included. Thus, the sign of the mitigation effect was stable, although its magnitude varied with the time since prior fire.

Together, these sensitivity analyses indicate that the main conclusions are robust to reasonable changes in threshold choices and prior-fire window definitions. The estimated magnitude of persistent loss and mitigation varies with threshold selection, but the qualitative results remain unchanged: persistent forest loss is widespread, crucial fire patterns are spatially heterogeneous rather than globally monotonic, fire severity remains central to crucial fire risk, and prior low-severity fire consistently reduces subsequent fire severity relative to matched controls.

#### Text S4. Field-based evaluation of MODIS post-fire recovery

To validate MODIS-derived post-fire recovery using independent field observations, we analyzed repeated live aboveground biomass measurements from the U.S. Forest Service Forest Inventory and Analysis (FIA) plots associated with documented fires[16]. FIA plot-fire events were assigned to WWF ecoregions, and biomass trajectories were expressed relative to their pre-fire values. We fitted hierarchical recovery models with a Gompertz growth model[3,17] and classified a site-event as recovered when its projected long-term biomass asymptote exceeded 95% of its pre-fire value. We estimated ecoregion recovery proportions using equal weights among eligible FIA plot-fire events. For comparison, MODIS recovery was defined using the same asymptotic threshold for TC and LAI, weighted by burned forest area, and summarized as the arithmetic mean of the two recovery proportions.

Eleven ecoregions had reportable FIA estimates, 19 had reportable MODIS estimates, and nine were jointly reportable (**Fig. S18**). Across the nine jointly reportable ecoregions, the 95% FIA and MODIS intervals overlapped in eight cases, but correspondence among the point estimates was weak (Pearson $r = -0.37$; RMSE = 0.18; normalized RMSE = 0.34; **Figs. S26-S27**). MODIS recovery proportions were, on average, 0.084 higher than FIA estimates. Thus, the FIA observations provided limited support for the broad magnitude of MODIS-derived recovery but did not demonstrate strong spatial agreement among ecoregions. This comparison should be interpreted cautiously because FIA represents plot-level live aboveground biomass and uses equal plot-fire-event weighting, whereas MODIS represents remotely sensed TC and LAI and uses burned-area weighting; their uncertainty intervals also quantify different sources of uncertainty.

#### Text S5. Relationships between fire-weather conditions and vegetation-based fire severity

We examined whether fire severity was associated with monthly Palmer Drought Severity Index (PDSI), precipitation, maximum temperature, and wind speed during the fire month. TerraClimate[18] values were linked to each valid fire event using its ignition year and month. Fire severity was summarized separately for TC, LAI, NPP, and ET as the event-level mean of non-negative first-year proportional vegetation loss. This event-level aggregation avoided treating multiple pixels within the same fire as independent observations. Relationships were evaluated using pooled correlations, equal-frequency binned trends, and rank associations calculated within region, biome, and fire-month strata (**Fig. S21**).

The pooled relationships were generally weak and inconsistent across vegetation indicators. Maximum temperature showed positive associations with TC and NPP severity, whereas wind speed showed negative associations for these metrics, but similar patterns were not evident for LAI and ET. Across all combinations, Spearman correlations ranged from -0.18 to 0.29, and linear models explained no more than 5.6% of the variation in severity. After accounting for broad differences among regions, biomes, and fire months, all rank associations remained small ($|r| \leq 0.10$), and their directions frequently differed among biomes. Thus, although individual weather–severity combinations showed localized patterns, no clear and consistent trend was observed between vegetation-based fire severity and the four monthly fire-weather indicators at the global scale.

#### Text S6. Uncertainty of mitigation effect

We quantified uncertainty in the fitted mitigation curves using 1,000 fire-cluster bootstrap replicates[19] because multiple matched pixels from the same fire event were not independent. Whole fire-event clusters were resampled with replacement, the same resampling frequency was applied to the control and prior-burn observations, and the generalized additive models were refitted for each vegetation indicator. Pointwise 95% uncertainty intervals were calculated from the 2.5% and 97.5% of the resulting curves. Low-severity prior-fire effects remained generally positive across much of the observed pre-fire range, particularly for TC, LAI, and NPP, whereas higher-severity prior-fire effects were commonly near zero or negative. Uncertainty increased near the ends of the observed vegetation ranges, where fewer fire events were available (**Extended Data Fig. 4**).

For the spatial analysis, each bootstrap realization of the biome-specific low-severity mitigation curves was propagated through the fixed global fire inventory and XGBoost recovery models[20]. For fires initially classified as crucial (recovery probability $p < 0.25$), counterfactual severity and recovery probability were recalculated, and a fire was considered mitigated when its probability increased to $p \geq 0.25$. Absolute spatial uncertainty was quantified as the standard deviation of annual mitigation density across curve realizations. Uncertainty was relatively low across most mapped hexagons but was higher in localized areas where variation in the mitigation curves more frequently changed whether crucial fires were classified as mitigated (**Extended Data Fig. 5**). Fire events, locations, and years were not resampled; thus, the mapped uncertainty represents uncertainty in the mitigation curves conditional on the observed fires and fixed recovery models, rather than interannual fire variability or XGBoost model-parameter uncertainty.

#### Supplementary tables

**Table S1. Model selection for postfire recovery trajectory fitting.** Performance comparison of candidate recovery models for tree cover (TC), leaf area index (LAI), net primary productivity (NPP), and evapotranspiration (ET). Models were evaluated using $R^2$, root mean square error (RMSE), and mean absolute error (MAE), with higher $R^2$ and lower RMSE and MAE indicating better performance. Bold values indicate the best-performing model for each vegetation indicator and evaluation criterion.

| | | Linear | Logistic | Monod | Chapman Richards | Negative exponent | Power | Gompertz |
|---|---|---|---|---|---|---|---|---|
| $R^2$ (↑) | TC | 0.572 | **0.608** | 0.460 | 0.552 | 0.504 | 0.587 | 0.604 |
| | LAI | 0.608 | **0.650** | 0.523 | 0.604 | 0.562 | 0.632 | 0.648 |
| | NPP | 0.606 | 0.656 | 0.589 | 0.643 | 0.612 | 0.634 | **0.679** |
| | ET | 0.599 | 0.650 | 0.551 | 0.623 | 0.580 | 0.622 | **0.662** |
| RMSE (↓) | TC | 0.344 | **0.332** | 0.403 | 0.366 | 0.384 | 0.339 | 0.337 |
| | LAI | 0.293 | **0.280** | 0.340 | 0.306 | 0.323 | 0.284 | 0.284 |
| | NPP | 0.193 | 0.179 | 0.204 | 0.186 | 0.195 | 0.186 | **0.172** |
| | ET | 0.223 | 0.207 | 0.247 | 0.221 | 0.235 | 0.217 | **0.204** |
| MAE (↓) | TC | 0.286 | **0.273** | 0.336 | 0.300 | 0.320 | 0.276 | **0.273** |
| | LAI | 0.241 | **0.228** | 0.279 | 0.248 | 0.265 | 0.231 | **0.228** |
| | NPP | 0.159 | 0.147 | 0.169 | 0.153 | 0.161 | 0.151 | **0.141** |
| | ET | 0.185 | 0.171 | 0.206 | 0.182 | 0.195 | 0.178 | **0.167** |

### Supplementary figures

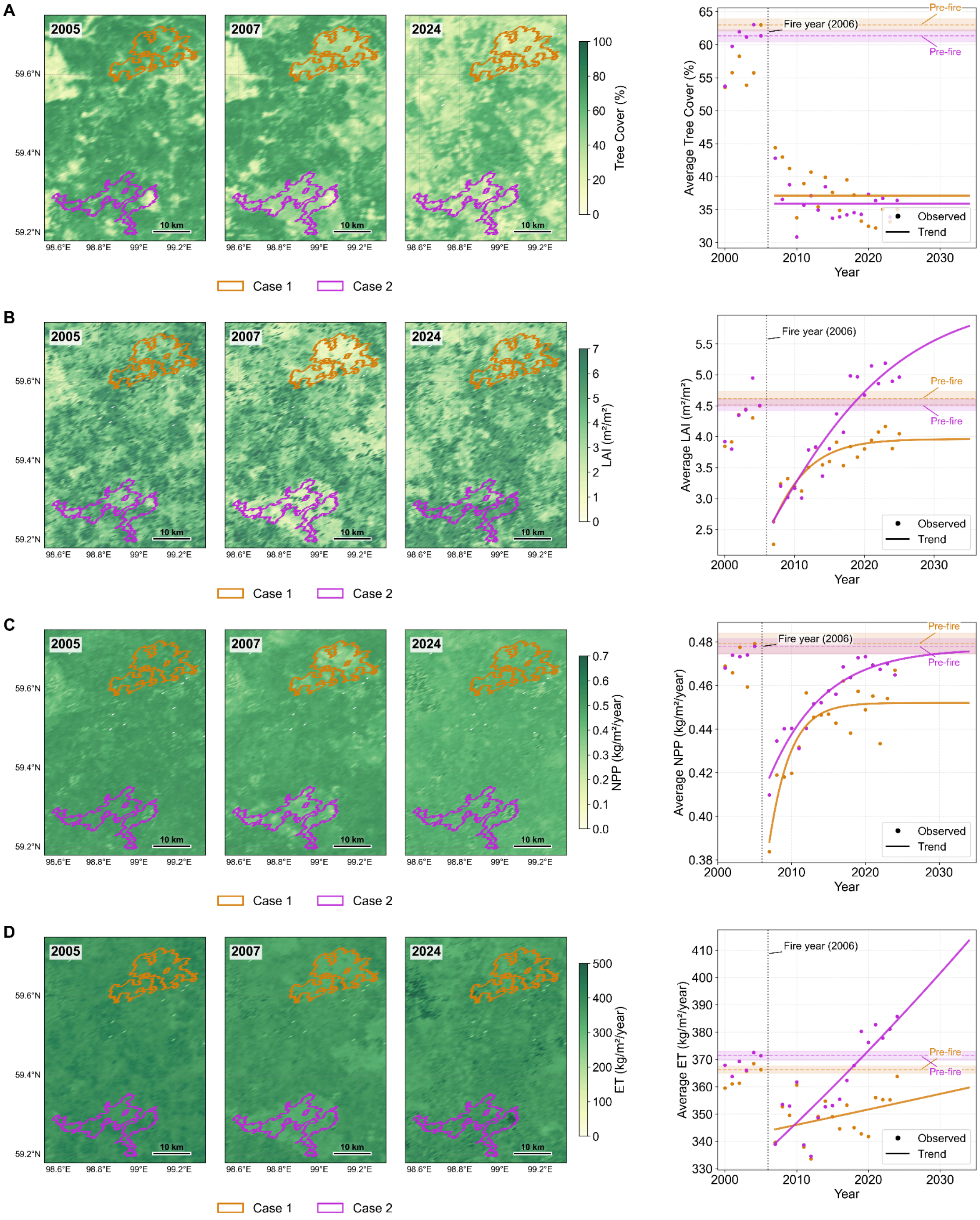


**Figure S1**. **Multi-indicator postfire trajectories for two contrasting Siberian forest cases.** (A–D) Spatial maps and temporal trajectories of tree cover (A), leaf area index (LAI; B), net primary productivity (NPP; C), and evapotranspiration (ET; D) for a boreal case-study region in Siberia affected by fire in 2006. Maps show conditions before the fire (2005), immediately after the fire (2007), and in 2024. Orange and magenta outlines denote a persistent-loss case and a recovered case, respectively. Right-hand panels show annual mean values within the two outlined cases, with points indicating observations and solid lines indicating fitted trends. Horizontal dashed lines and shaded bands indicate the prefire baseline and its uncertainty range for each case, and the vertical dotted line marks the 2006 fire year.

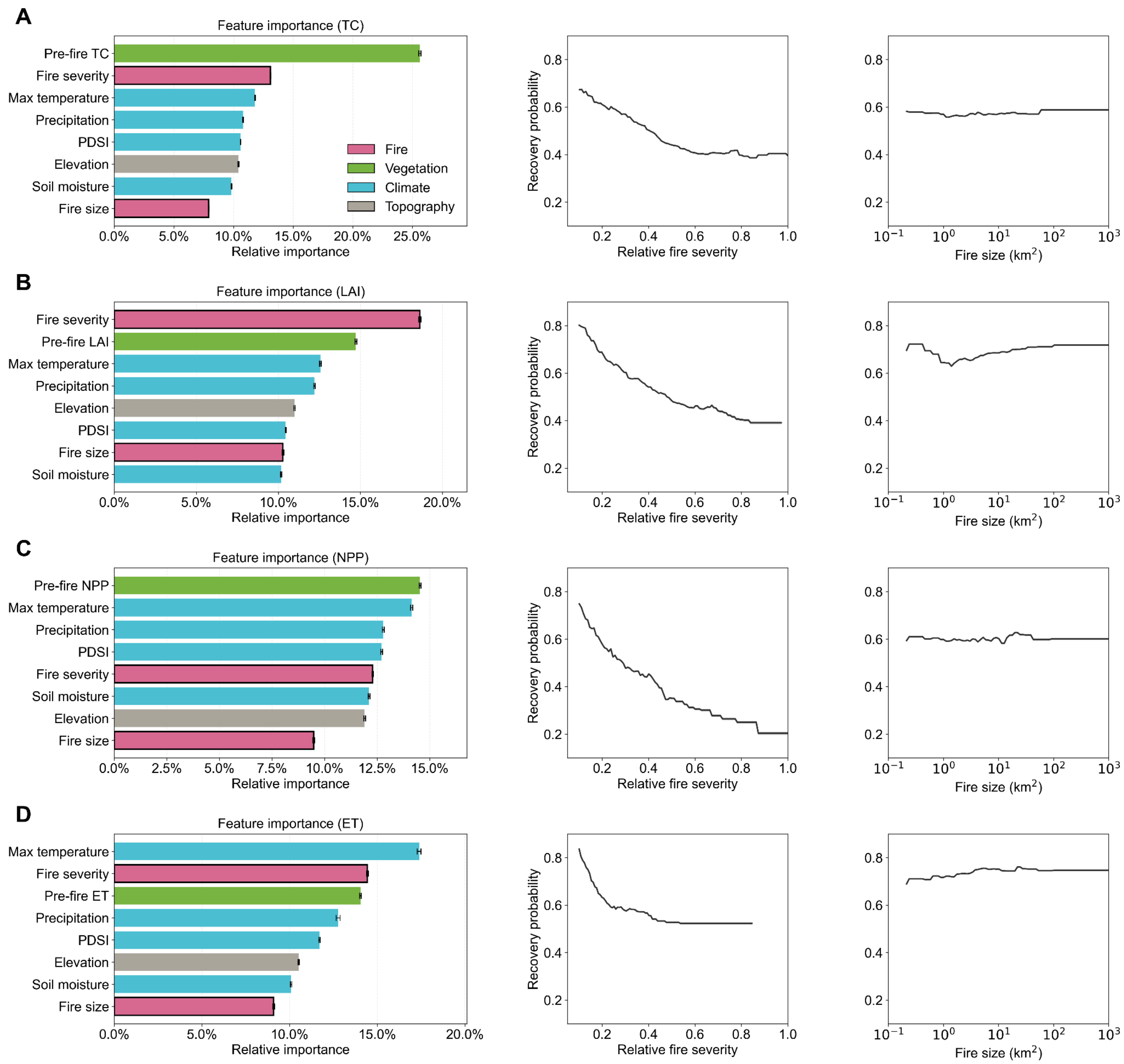


**Figure S2**. **XGBoost importance and PDP curve.** (A–D) The recovery probability model (XGBoost) of tree cover (A), leaf area index (LAI; B), net primary productivity (NPP; C), and evapotranspiration (ET; D) , feature-importance scores identify fire severity, pre-fire vegetation indicators, and climate variables as the main predictors of recovery probability; partial dependence plots show declining recovery probability with increasing fire severity and relatively stable probability with fire size change.

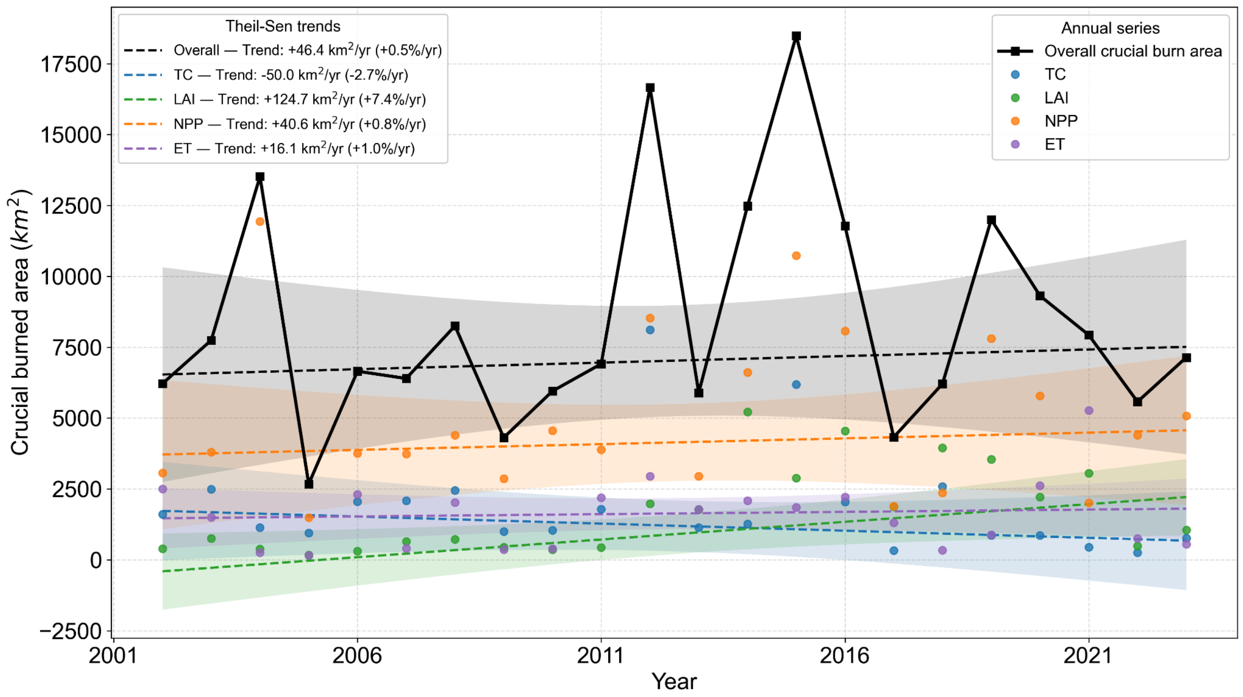


**Figure S3**. **Global temporal trends in crucial burned area.** Annual global crucial burned area from 2001 to 2023, summarized for the combined estimate and for individual structural and functional indicators, including tree cover (TC), leaf area index (LAI), net primary productivity (NPP), and evapotranspiration (ET). Points indicate annual indicator-specific estimates, dashed lines show fitted temporal trends, and shaded bands indicate trend uncertainty.

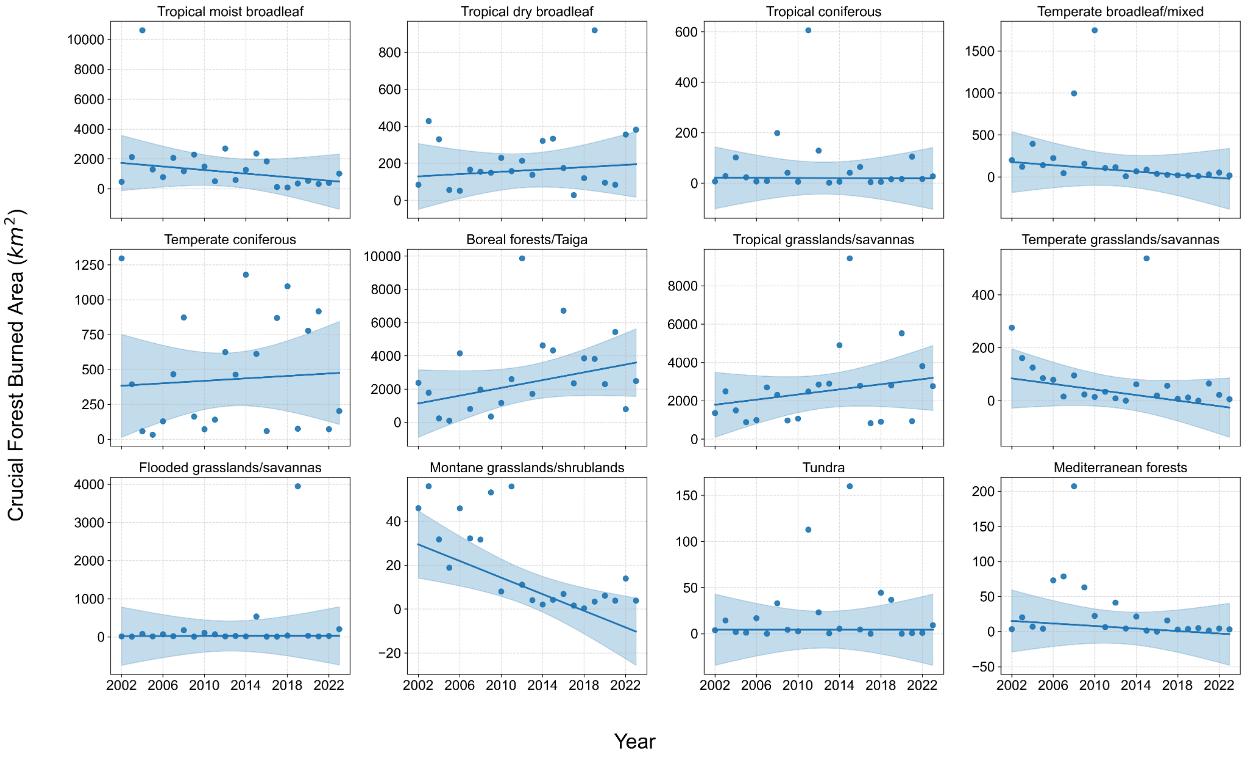

**Figure S4**. **Biome-specific temporal trends in crucial forest burned area.** Annual crucial forest burned area from 2001 to 2023 across major global biomes. Each panel represents one biome, with points indicating annual estimates, solid lines showing fitted temporal trends, and shaded bands indicating trend uncertainty.

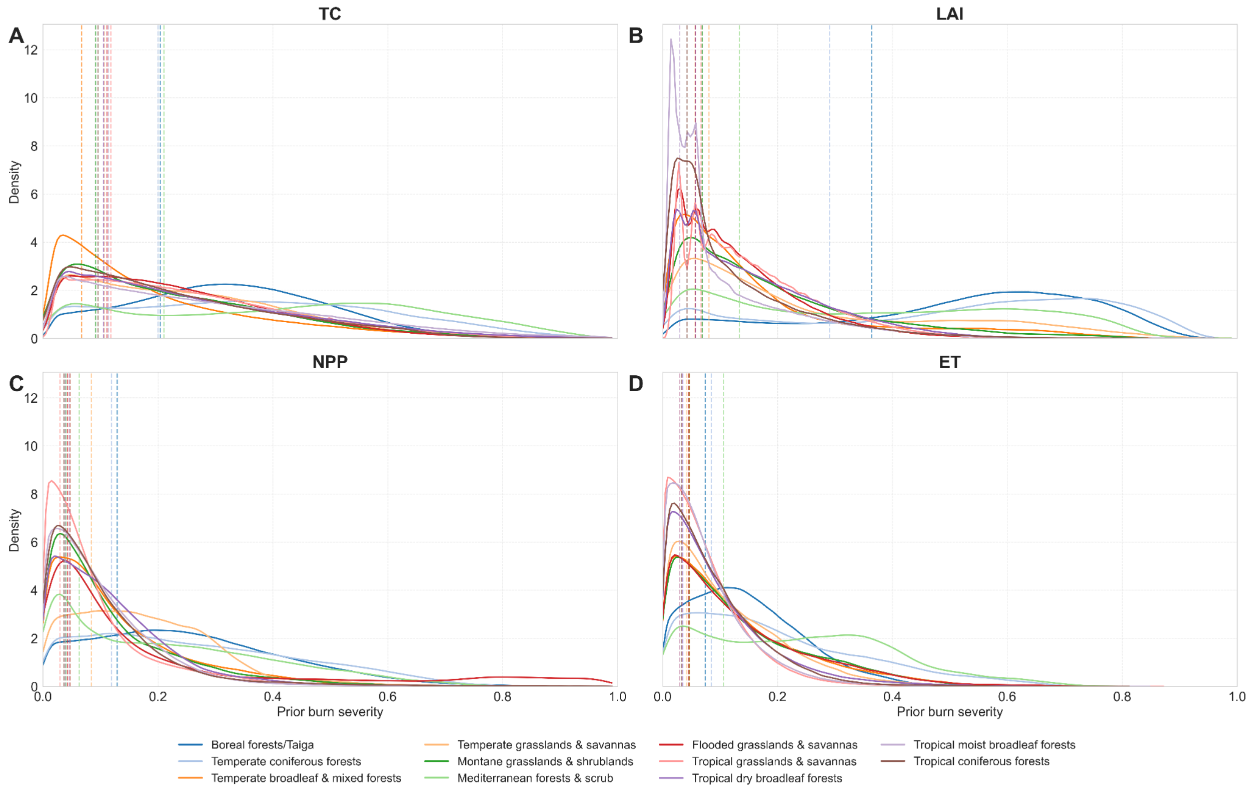


**Figure S5**. **Biome-specific distributions of prior burn severity.**Density distributions of prior burn severity across major global biomes, calculated separately for tree cover (TC; A), leaf area index (LAI; B), net primary productivity (NPP; C), and evapotranspiration (ET; D). Colored curves represent biome-specific severity distributions. Vertical dashed lines mark the biome-specific severity value corresponding to the 25% cumulative level of the summed density, indicating the lower-severity portion of each distribution.

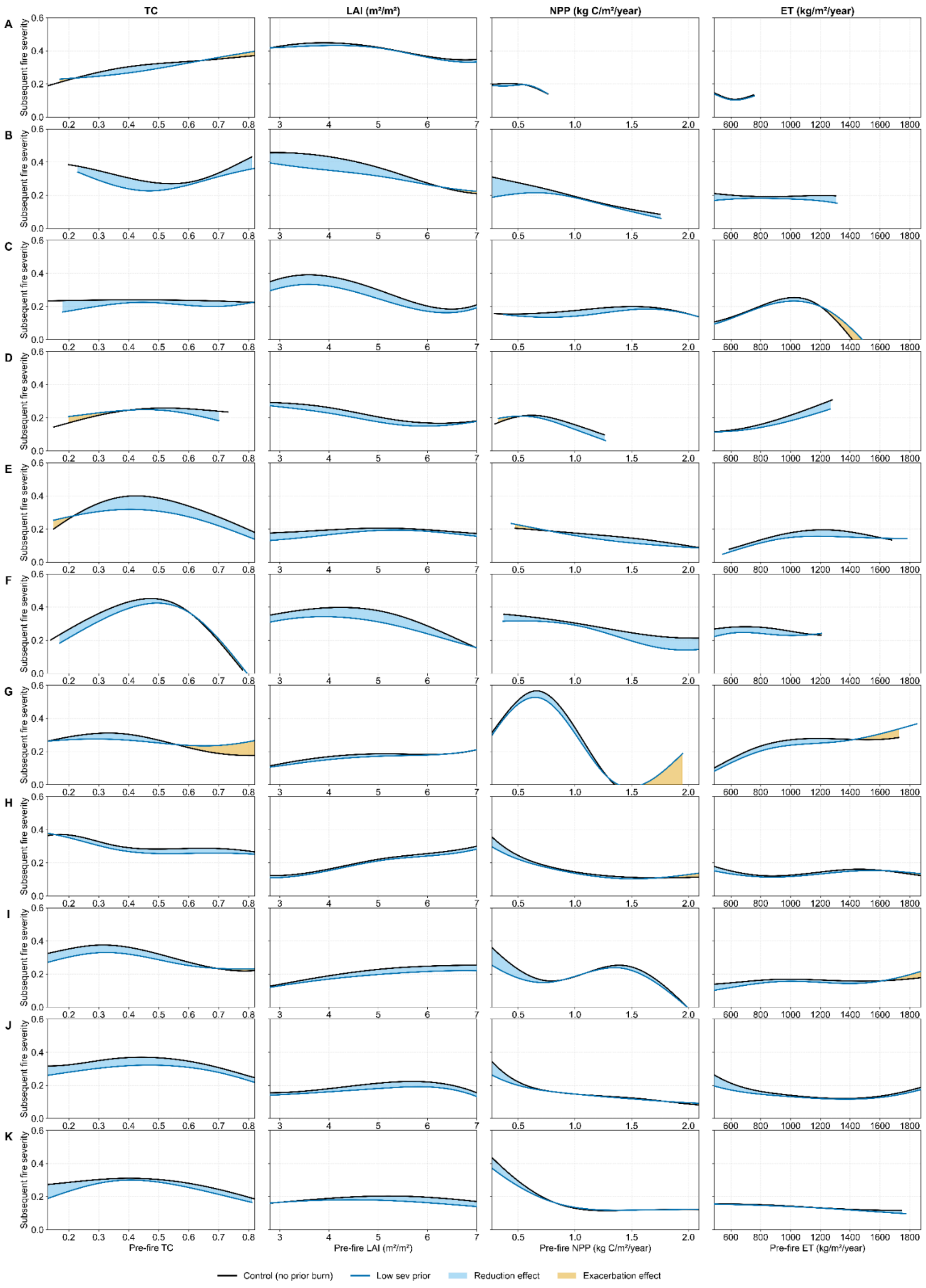


**Figure S6**. **Biome-specific effects of low-severity prior burns on subsequent fire severity.** Relationships between prefire ecosystem condition and new fire severity for areas with low-severity prior burns compared with matched control areas without prior burn. Columns show structural and functional indicators, including tree cover (TC), leaf area index (LAI), net primary productivity (NPP), and evapotranspiration (ET). Rows represent major biomes: boreal forests/taiga (A), temperate coniferous forests (B), temperate broadleaf and mixed forests (C), temperate grasslands and savannas (D), montane grasslands and shrublands (E), Mediterranean forests and scrub (F), flooded grasslands and savannas (G), tropical grasslands and savannas (H), tropical dry broadleaf forests (I), tropical moist broadleaf forests (J), and tropical coniferous forests (K). Black and blue curves indicate fitted responses for control and low-severity prior-burn areas, respectively. Blue shading indicates reduced subsequent fire severity relative to controls, whereas yellow shading indicates exacerbated severity.

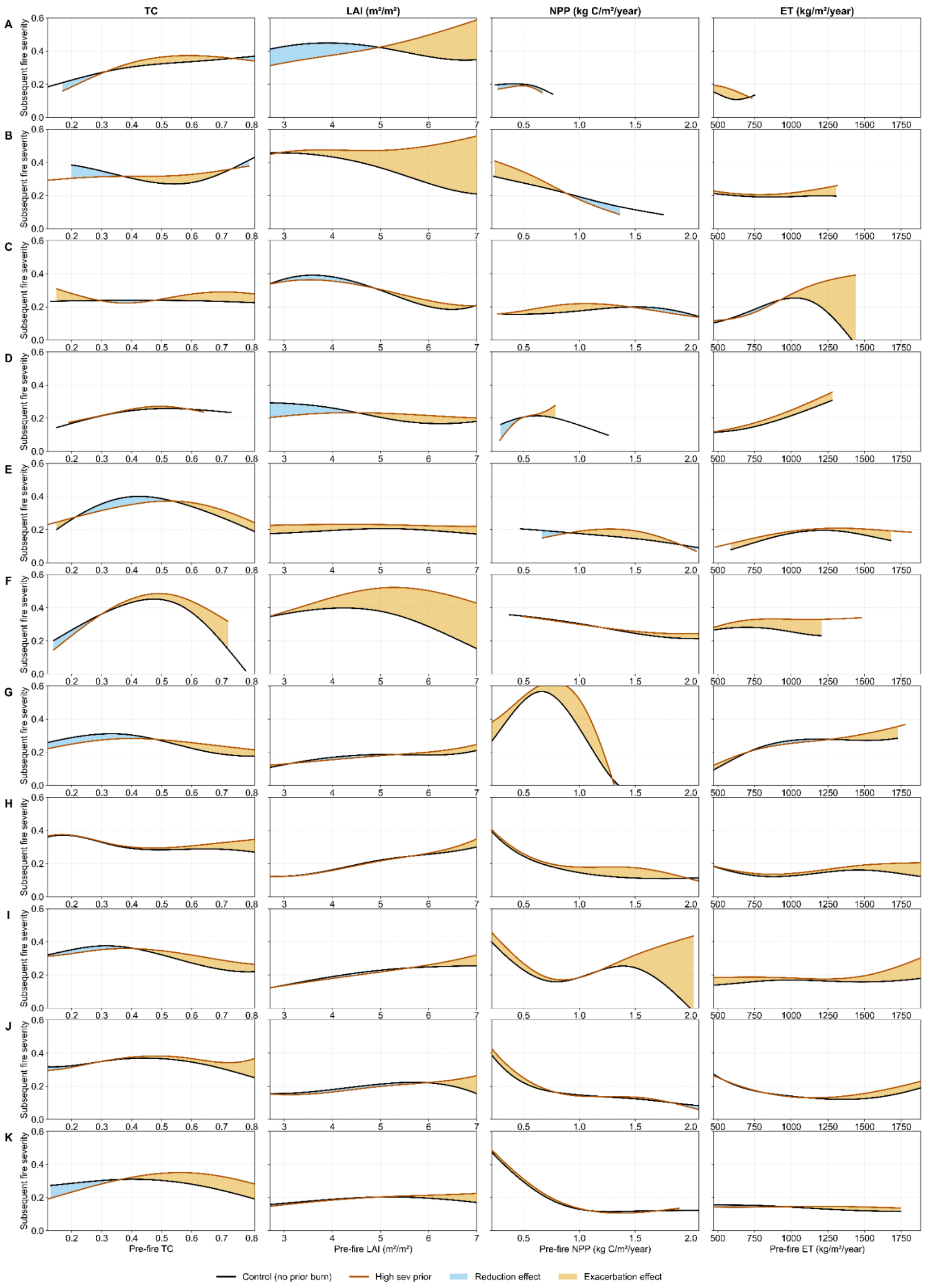


**Figure S7**. **Biome-specific effects of high-severity prior burns on subsequent fire severity.** Relationships between prefire ecosystem condition and new fire severity for areas with high-severity prior burns compared with matched control areas without prior burn. Columns show TC, LAI, NPP, and ET, and rows follow the same biome order as in Fig. S6. Rows represent major biomes: boreal forests/taiga (A), temperate coniferous forests (B), temperate broadleaf and mixed forests (C), temperate grasslands and savannas (D), montane grasslands and shrublands (E), Mediterranean forests and scrub (F), flooded grasslands and savannas (G), tropical grasslands and savannas (H), tropical dry broadleaf forests (I), tropical moist broadleaf forests (J), and tropical coniferous forests (K). Black and blue curves indicate fitted responses for control and low-severity prior-burn areas, respectively. Blue shading indicates reduced subsequent fire severity relative to controls, whereas yellow shading indicates exacerbated severity. Black and orange curves indicate fitted responses for control and high-severity prior-burn areas, respectively. Blue shading indicates a reduction effect, where prior burns are associated with lower subsequent fire severity than controls, whereas yellow shading indicates an exacerbation effect, where prior burns are associated with higher subsequent fire severity.

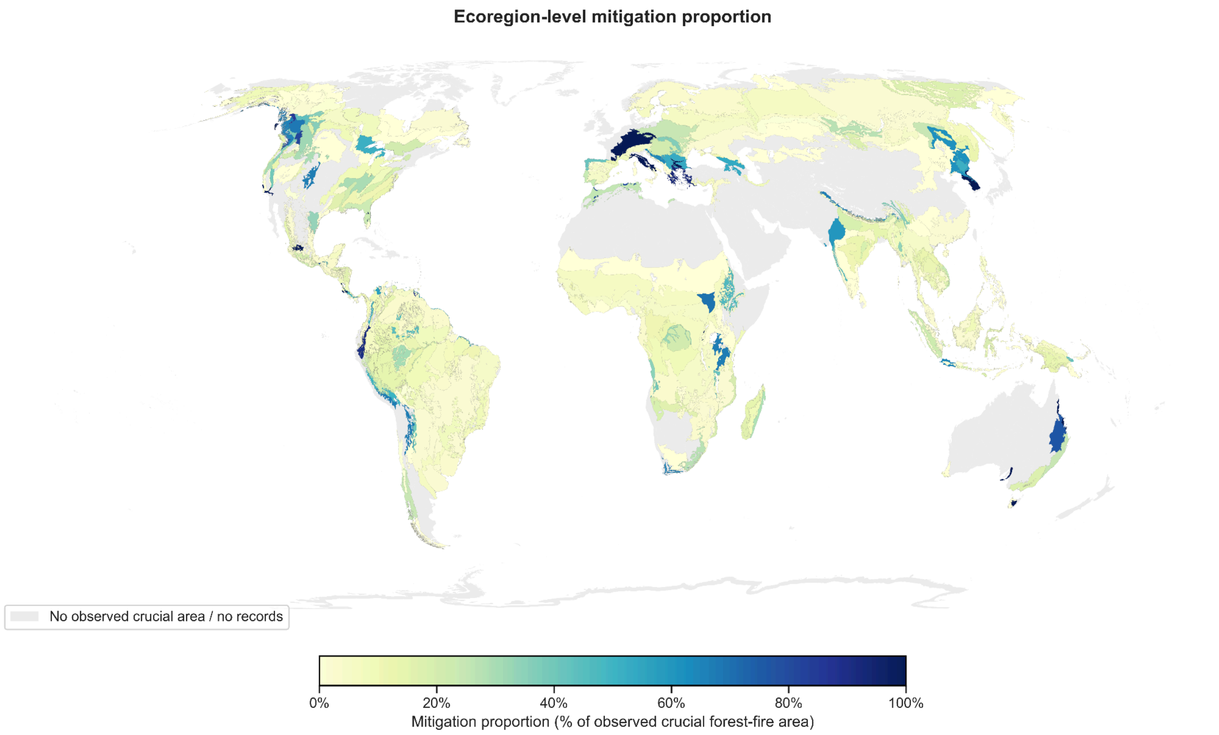


**Figure S8**. **Ecoregion-level mitigation proportion under a low-severity prior-burn scenario.** Spatial distribution of mitigation proportion across global ecoregions, expressed as the percentage of observed crucial forest-fire area that could be mitigated under the low-severity prior-burn scenario. Darker colors indicate ecoregions with higher relative mitigation potential, whereas pale colors indicate limited mitigation potential. Gray areas denote ecoregions with no observed crucial forest-fire area or insufficient fire records.

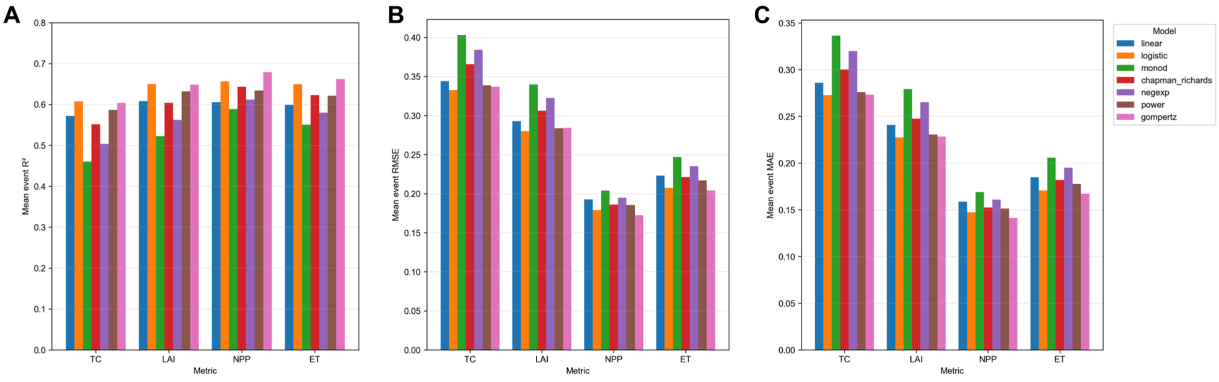


**Figure S9. Summary of candidate model performance for postfire recovery fitting.** Comparison of seven candidate recovery models across four vegetation indicators including TC, LAI, NPP, and ET. Panels show event-level mean $R^2$ (A), RMSE (B), and MAE (C). Higher $R^2$ and lower RMSE and MAE indicate better model performance.

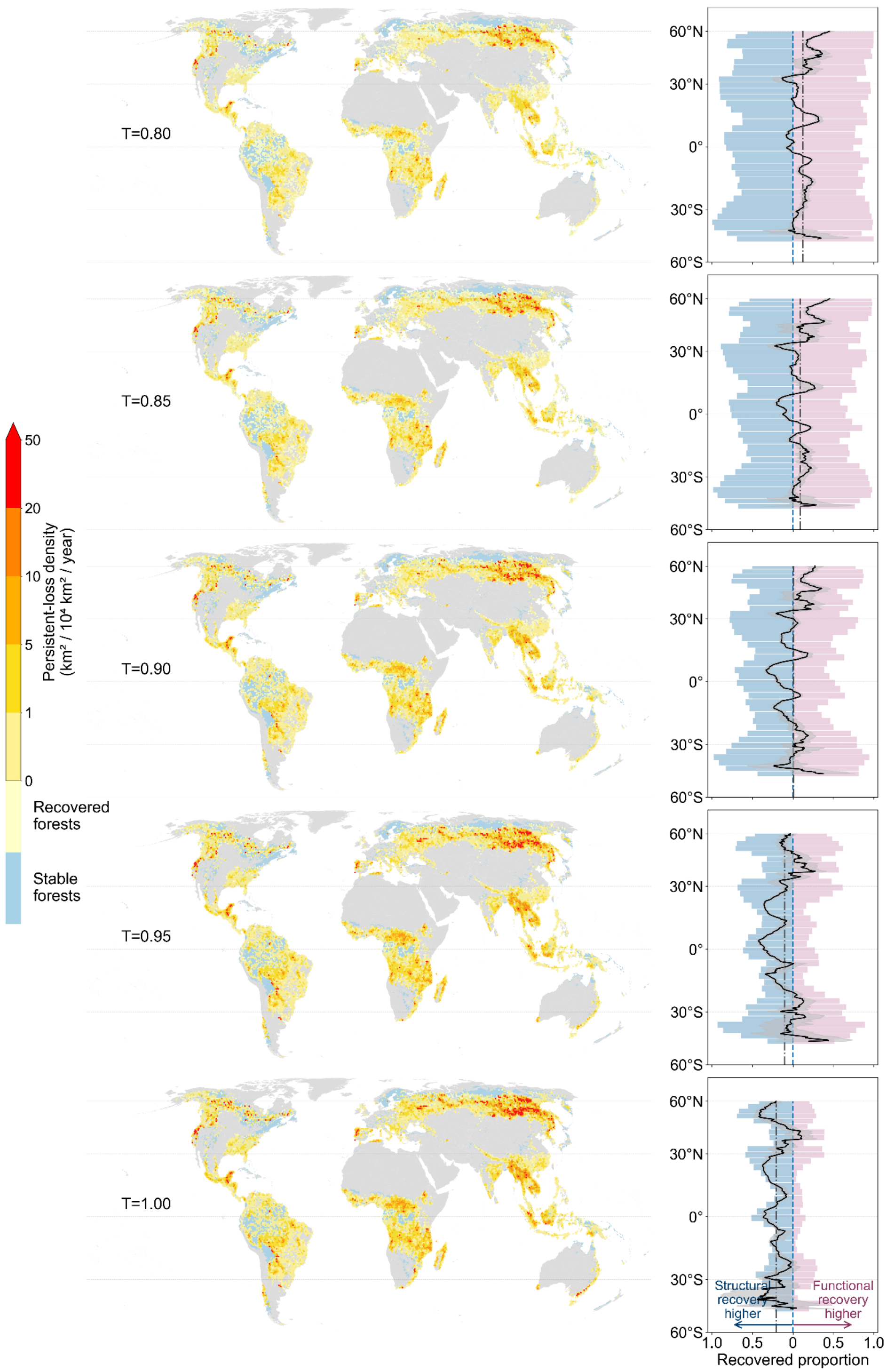


**Figure S10. Sensitivity of persistent-loss spatial patterns to recovery-potential thresholds.** Global persistent-loss density under alternative recovery-potential thresholds from 0.80 to 1.00. Left panels show persistent-loss density on 50-km equal-area hexagons, with warmer colors indicating higher annual persistent-loss density, pale yellow indicating recovered forests, and blue indicating stable forests without recorded fire. Right panels show the corresponding latitudinal profiles of recovered proportion for structural indicators (TC and LAI) and functional indicators (NPP and ET). Blue and pink bars indicate structural and functional recovered proportions, respectively. The black curve shows the structural-functional recovery contrast with uncertainty shading. The blue dashed vertical line marks zero contrast, and the gray dash-dot vertical line marks the mean structural-functional recovery contrast across latitudes for each threshold.

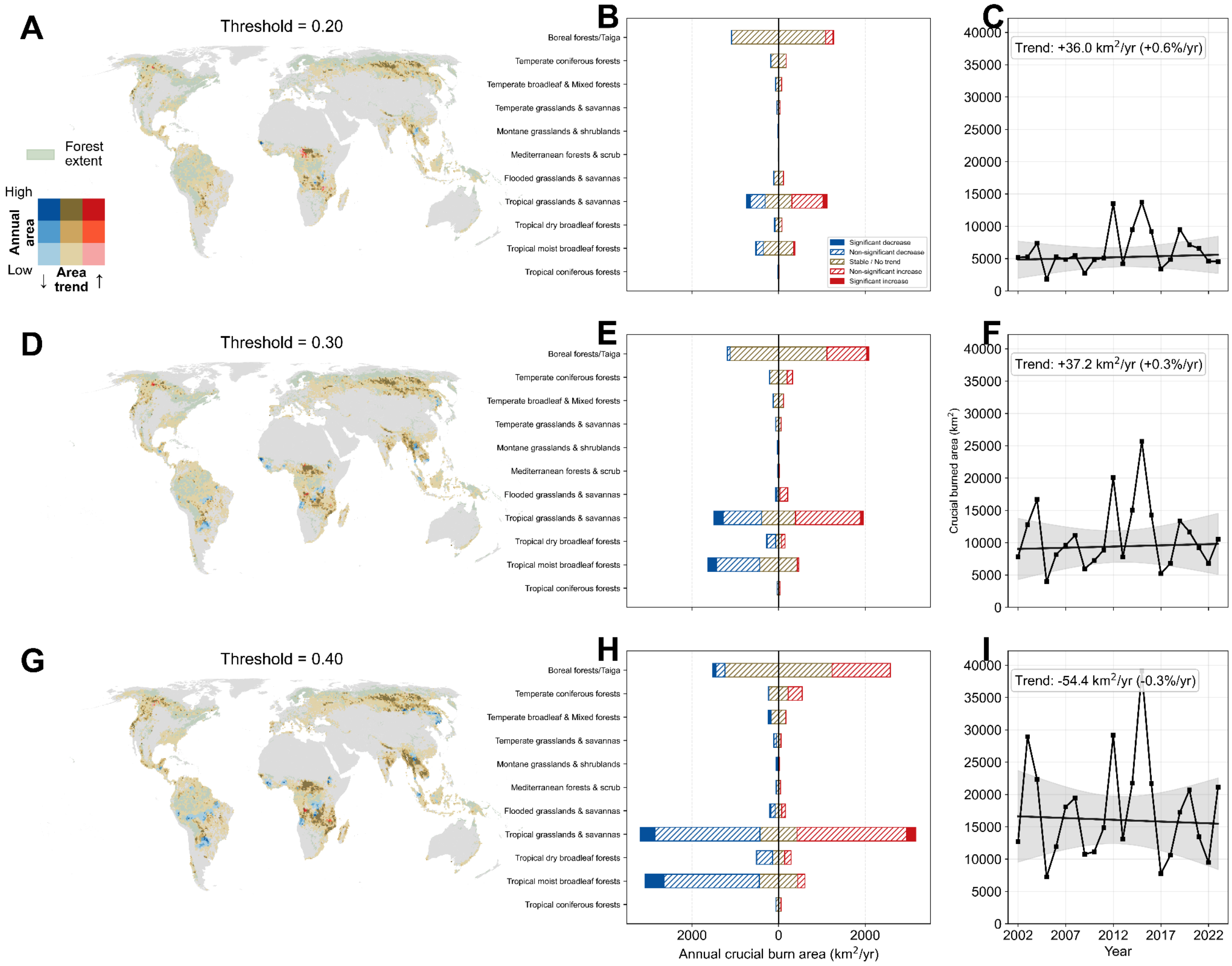


**Figure S11. Spatial and temporal sensitivity of crucial fire estimates to alternative probability thresholds.** Spatial, biome-level, and temporal patterns of crucial fires under probability thresholds of 0.20, 0.25, and 0.30. Rows correspond to threshold values of 0.20 (A–C), 0.30 (D–F), and 0.40 (G–I). (A, D, G) Global spatial patterns of crucial fires, with colors jointly representing annual crucial burned area and temporal trend direction, and pale green indicating forest extent. (B, E, H) Biome-level annual crucial burned area partitioned by trend direction and statistical significance. (C, F, I) Global annual crucial burned area from 2002 to 2023, with fitted trends, uncertainty bands, and inset values reporting trend slopes.

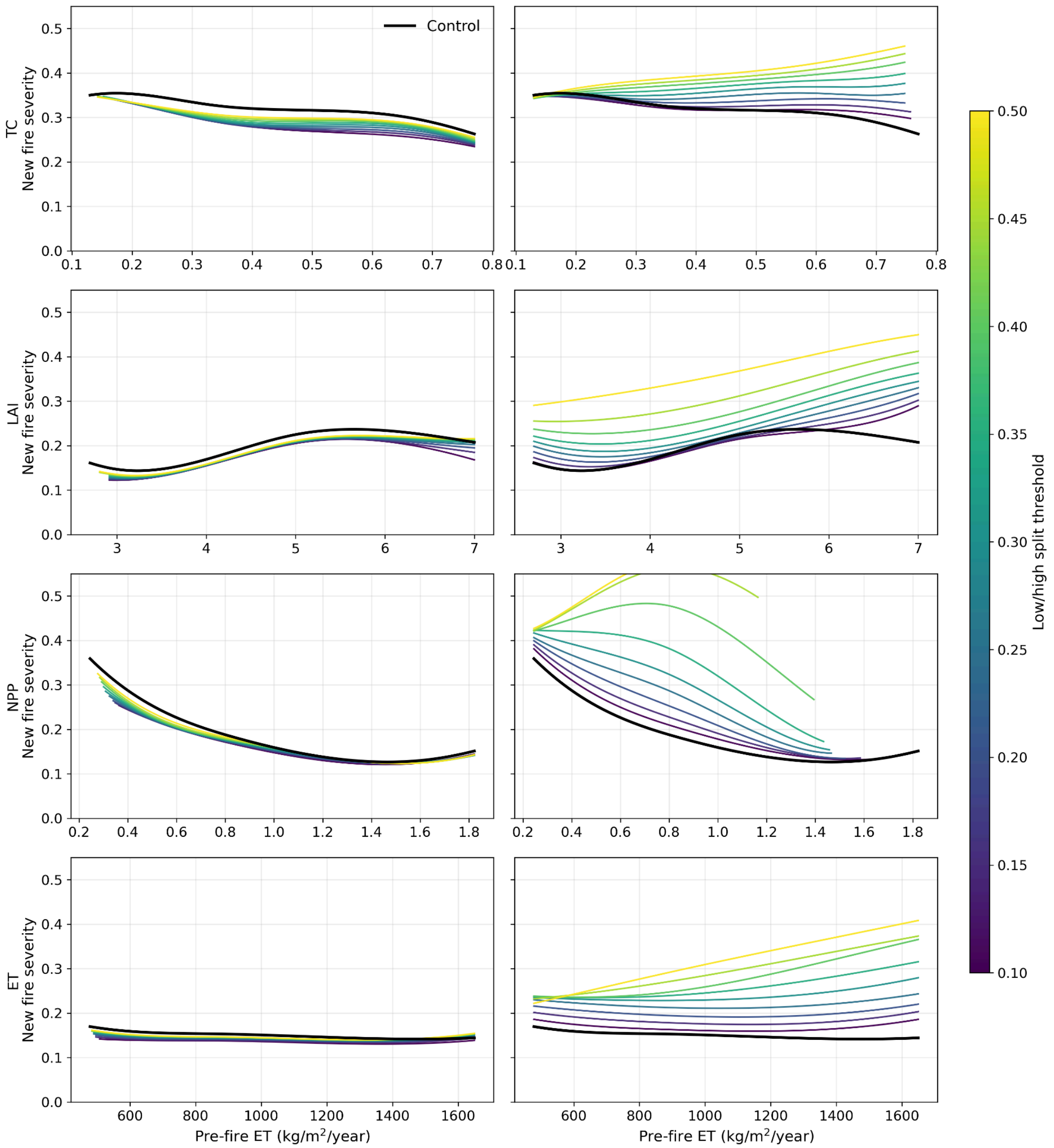


**Figure S12. Sensitivity of prior-burn effects to the low-severity threshold.** Fitted relationships between prefire vegetation condition and subsequent fire severity under alternative thresholds used to separate low- and high-severity prior burns. Rows show four vegetation indicators. The left column shows low-severity prior-burn mitigation effects, and the right column shows high-severity prior-burn effects. Colored curves indicate different threshold values, and black curves indicate controls without prior burn.

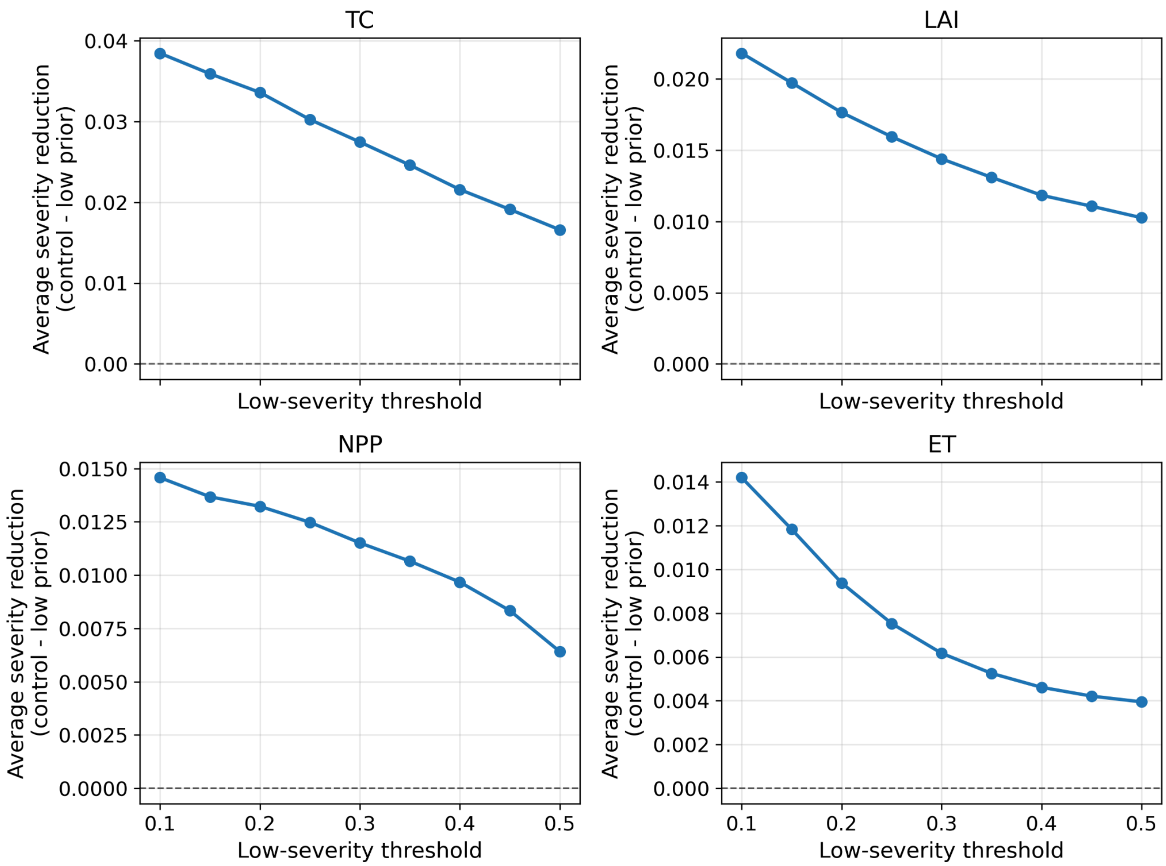


**Figure S13. Sensitivity of estimated mitigation effects to the low-severity threshold.** Average fire-severity reduction associated with low-severity prior burns was recalculated across low-severity thresholds from 0.10 to 0.50 for TC, LAI, NPP, and ET. The dashed horizontal line marks zero effect.

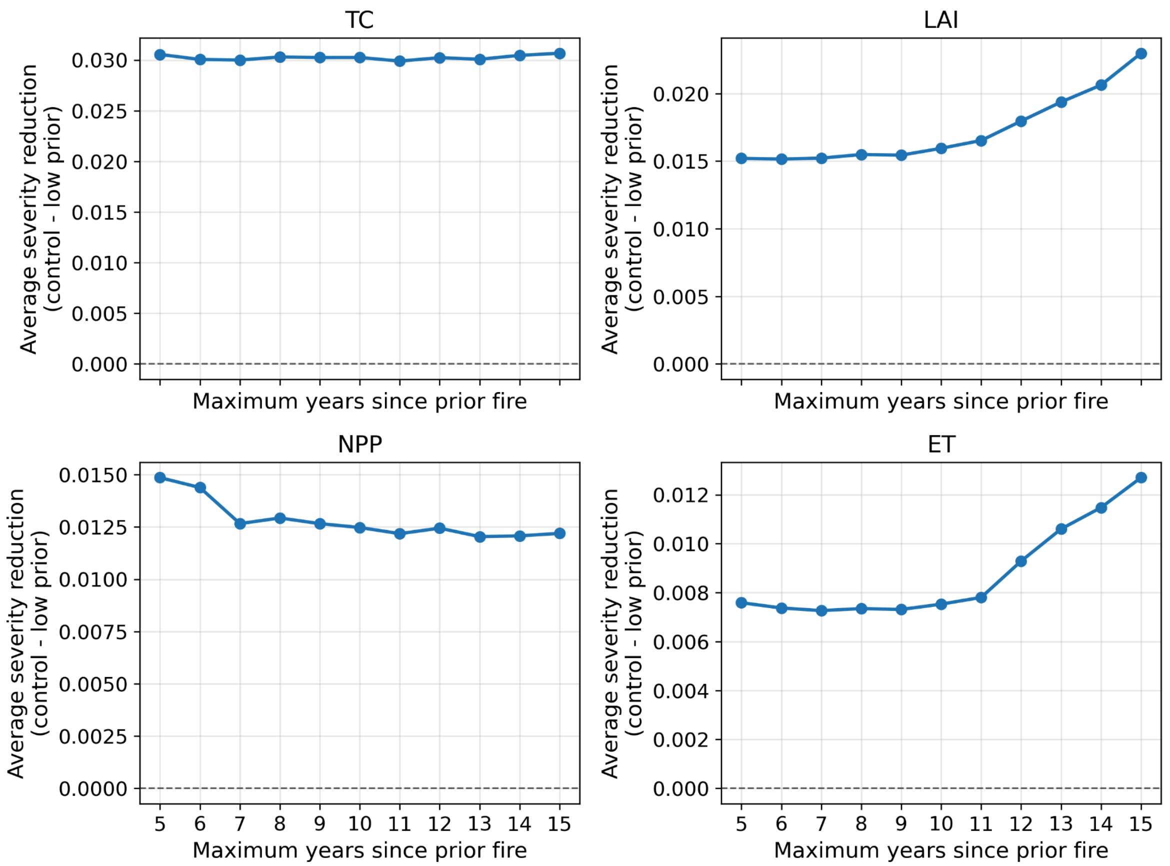


**Figure S14. Sensitivity of estimated mitigation effects to the prior-burn time window.** Average fire-severity reduction associated with low-severity prior burns was recalculated using maximum prior-fire windows from 5 to 15 years for TC, LAI, NPP, and ET. The dashed horizontal line marks zero effect.

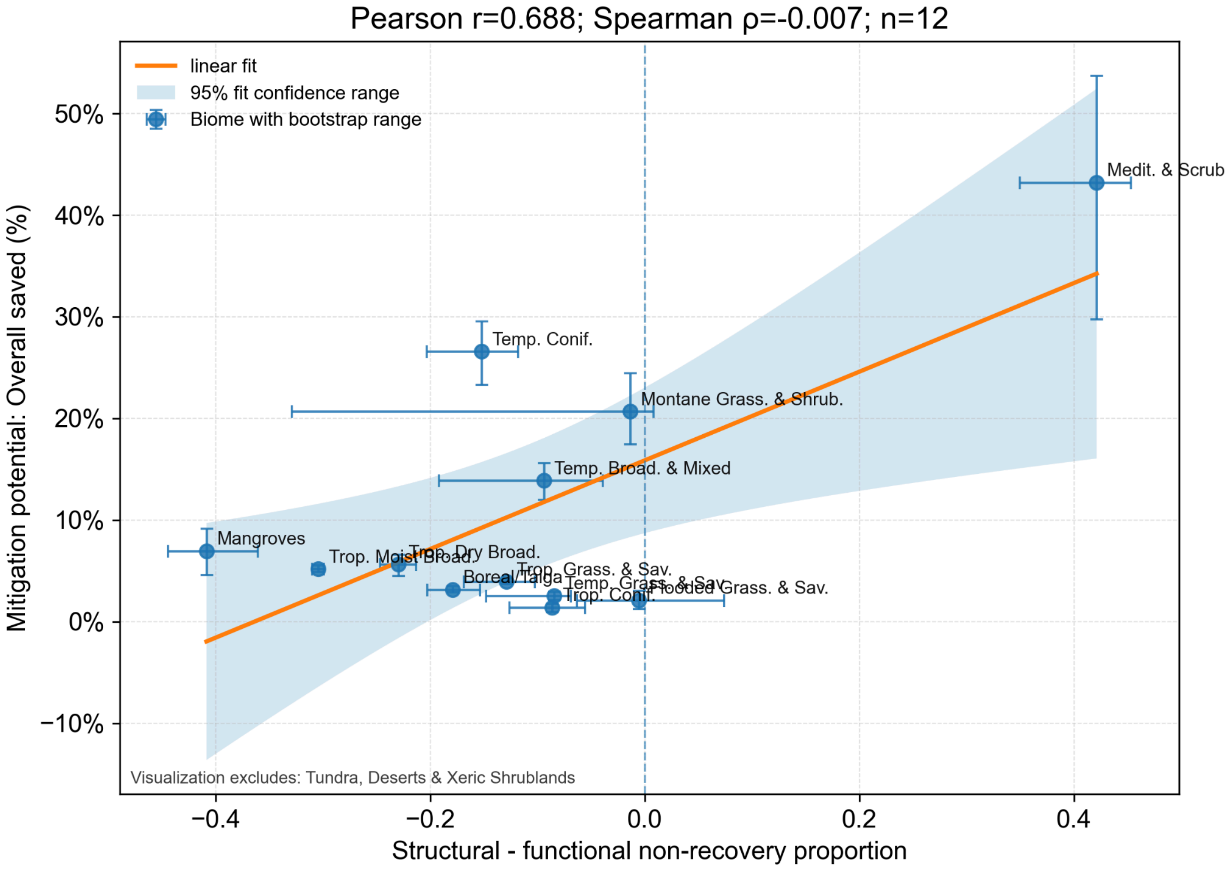


**Figure S15. Relationship between structural-functional recovery contrast and biome-level mitigation potential.** Biome-level mitigation potential was compared with the difference in recovery proportion between structural and functional metrics. Positive x-axis values indicate higher structural than functional recovery proportion, whereas negative values indicate higher functional recovery proportion. Points represent biome-level estimates, error bars indicate bootstrap ranges from 25% - 75%, and the orange line and shaded band show the fitted linear relationship and 95% confidence interval.

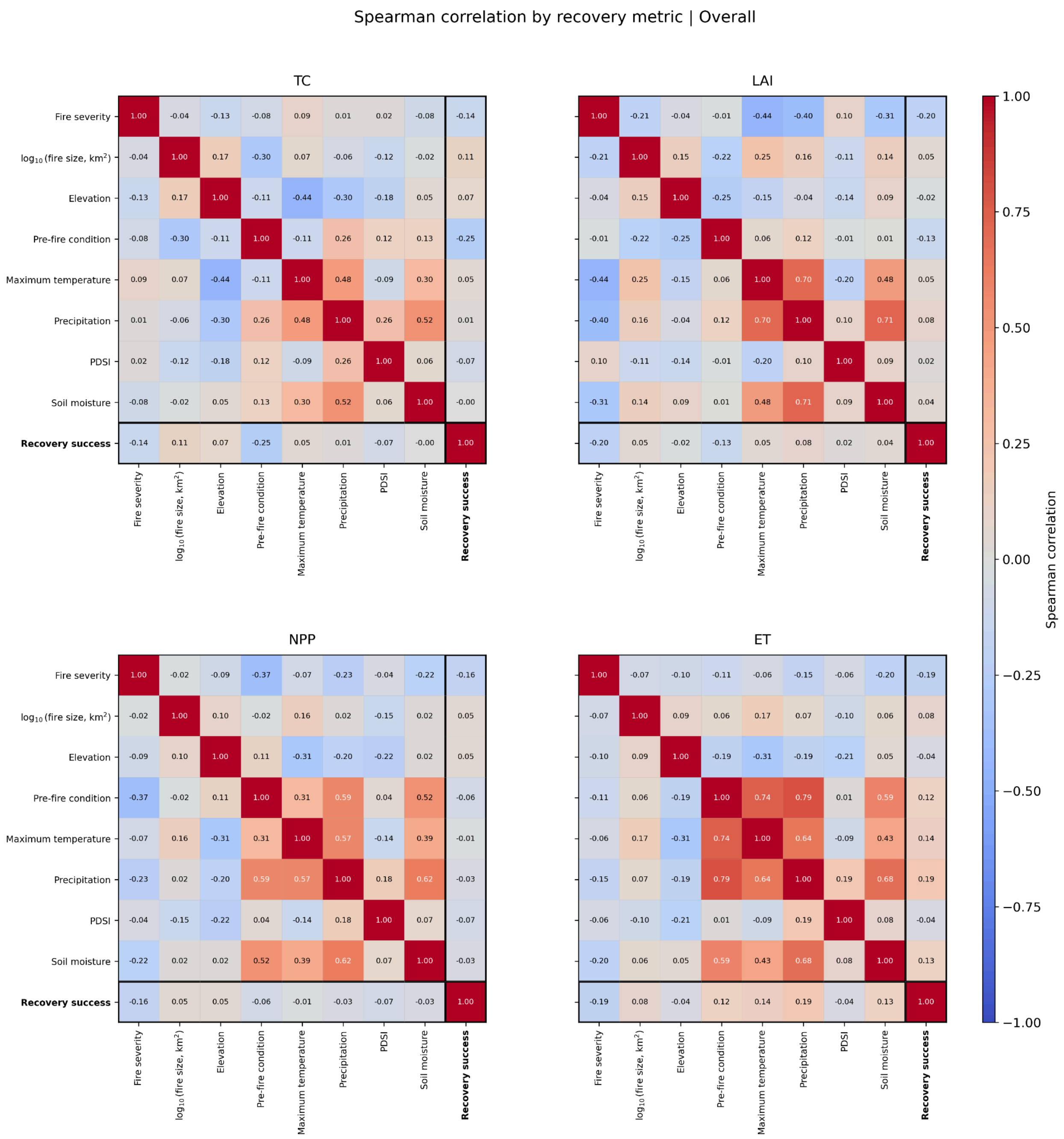


**Figure S16 Pairwise correlations among fire, environmental, and recovery variables.** Heatmaps show overall Spearman rank-correlation coefficients (ρ) for tree cover (TC), leaf area index (LAI), net primary productivity (NPP), and evapotranspiration (ET), pooled across seven regional datasets. Recovery success was defined as a metric-specific fitted recovery asymptote ≥0.95. Variables include fire severity, $\log_{10}$-transformed fire size, elevation, pre-fire condition, maximum temperature, precipitation, Palmer Drought Severity Index (PDSI), soil moisture, and recovery success. Cell values report ρ; red and blue indicate positive and negative correlations, respectively, with darker colors representing stronger relationships. Black outlines identify correlations involving recovery success, which were generally weak ($|\rho| \leq 0.25$).

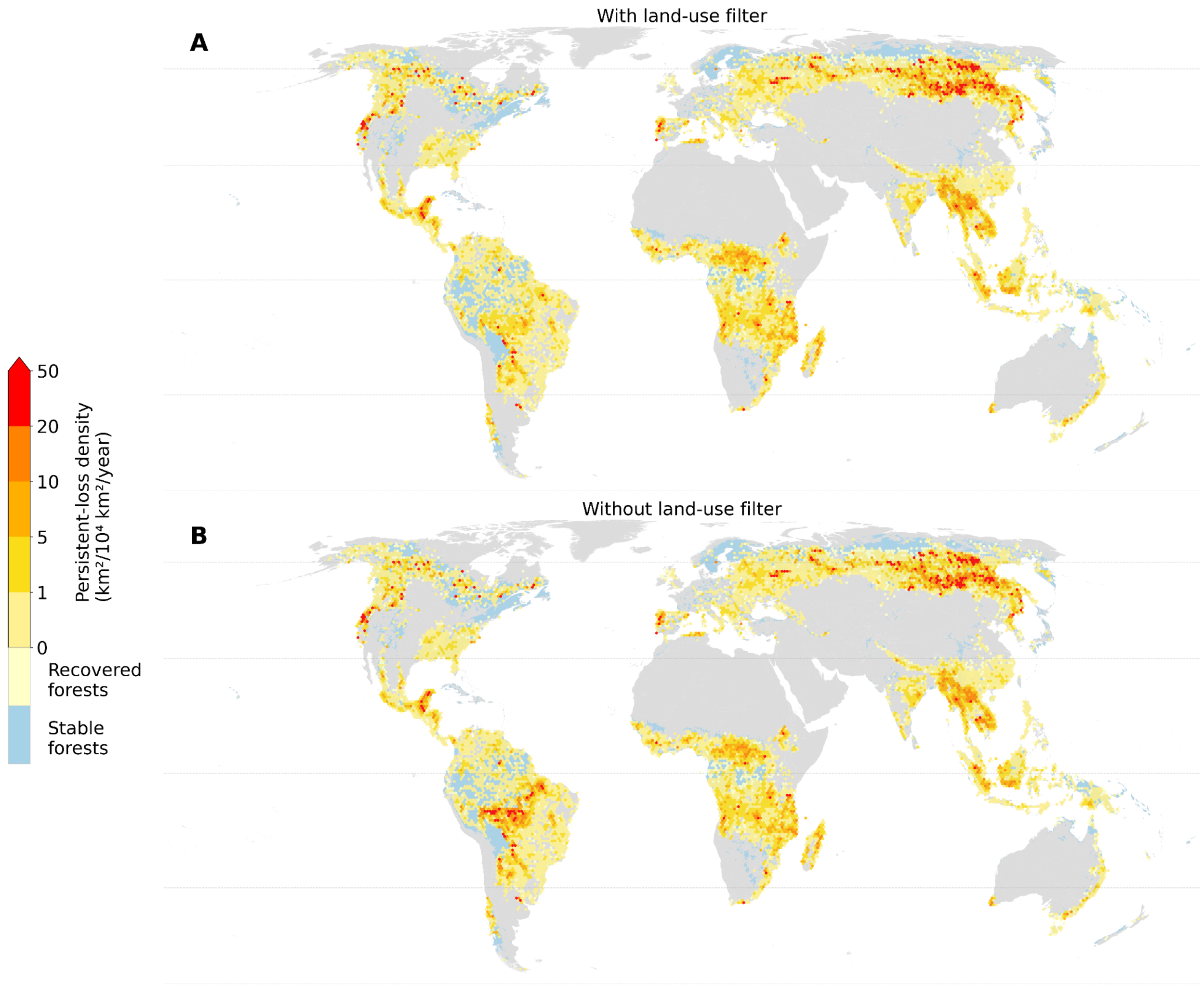


**Figure S17. Sensitivity of global persistent-loss patterns to land-use filtering.** Global persistent forest-loss density estimated with (A) and without (B) the land-use filter used to exclude fire events potentially affected by postfire land conversion. Maps are summarized on 50-km equal-area hexagons. Warmer colors indicate higher annual persistent-loss density, pale yellow indicates recovered forests, and blue indicates stable forests without recorded fire.

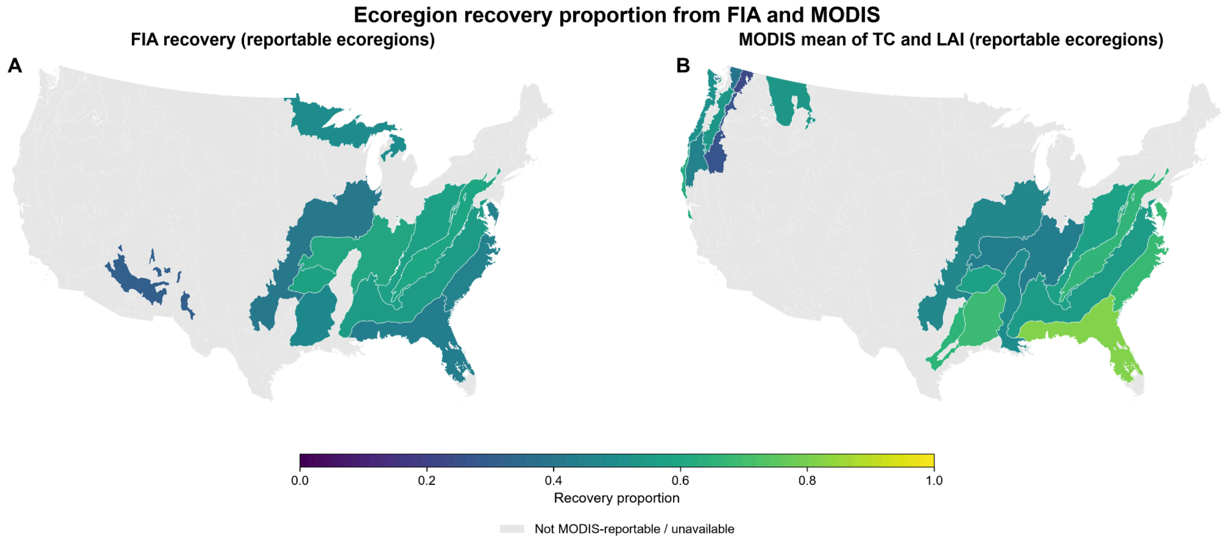


**Figure S18. Spatial distribution of FIA- and MODIS-based post-fire recovery proportions across WWF ecoregions in the conterminous United States.** (A) FIA recovery proportions for 11 reportable ecoregions. (B) Arithmetic mean of the MODIS TC- and LAI-based recovery proportions for 19 MODIS-reportable ecoregions. Both panels use the same recovery-proportion scale, with higher values indicating a greater proportion of events projected to recover above 95% of their pre-fire condition. Gray indicates ecoregions without an available or reportable estimate.

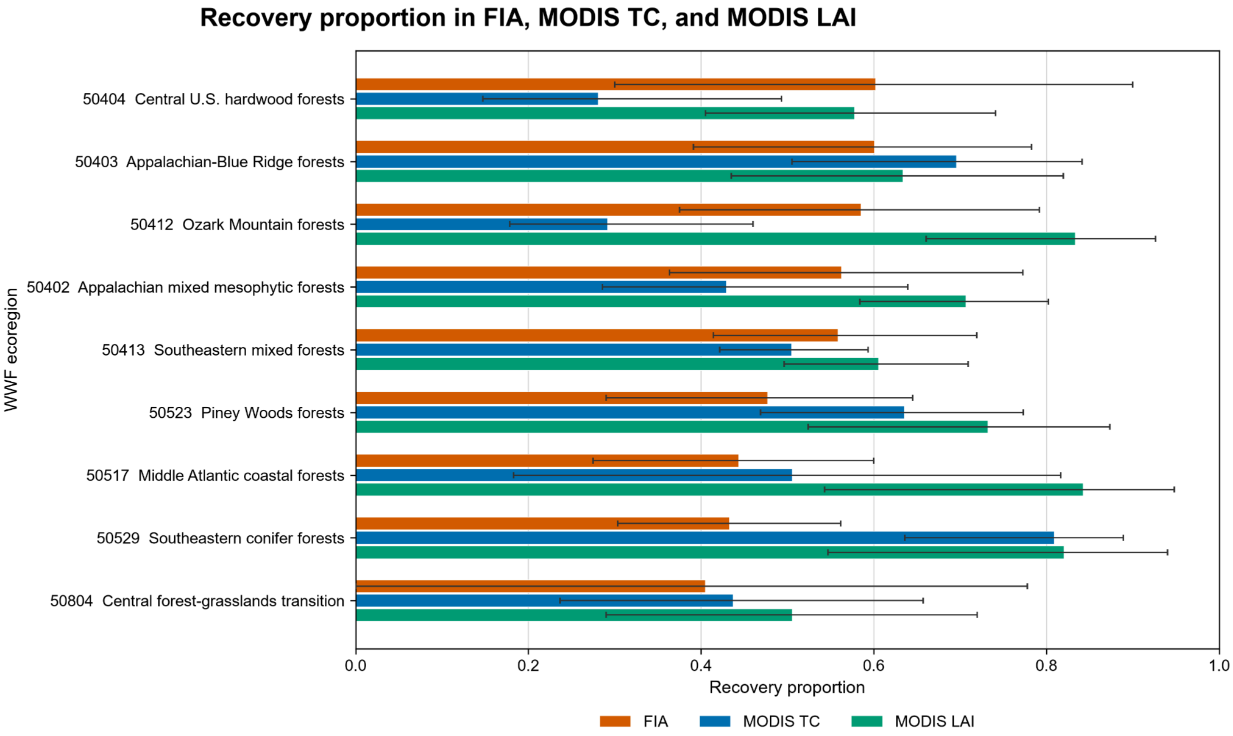


**Figure S19. Comparison of FIA- and MODIS-based post-fire recovery proportions across nine jointly reportable WWF ecoregions.** Orange bars show FIA estimates based on equal weighting of eligible plot–fire events, whereas blue and green bars show burned-forest-area-weighted MODIS tree cover (TC) and leaf area index (LAI) estimates, respectively. Recovery was defined as a fitted long-term asymptote exceeding 95% of the pre-fire value. Error bars show 95% uncertainty intervals derived from the FIA hierarchical analysis or the MODIS fire-event cluster bootstrap.

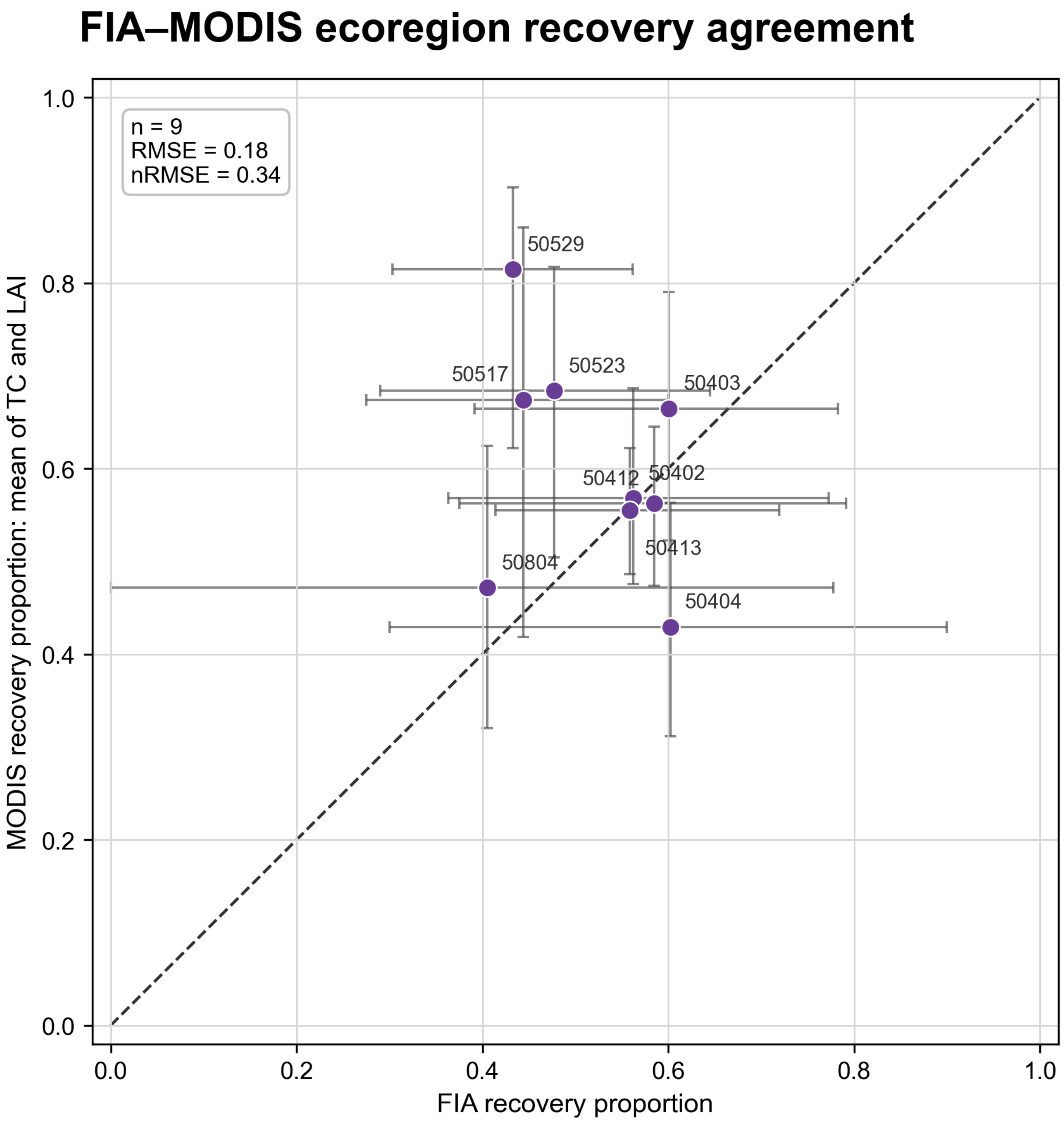


**Figure S20. Agreement between FIA- and MODIS-based recovery proportions across nine jointly reportable WWF ecoregions.** Each point compares the FIA recovery proportion with the arithmetic mean of the MODIS TC and LAI recovery proportions. Horizontal and vertical error bars show the respective 95% uncertainty intervals, and the dashed line indicates 1:1 agreement. The root mean square error (RMSE) was 0.18, and the normalized RMSE was 0.34, calculated as RMSE divided by the mean FIA recovery proportion.

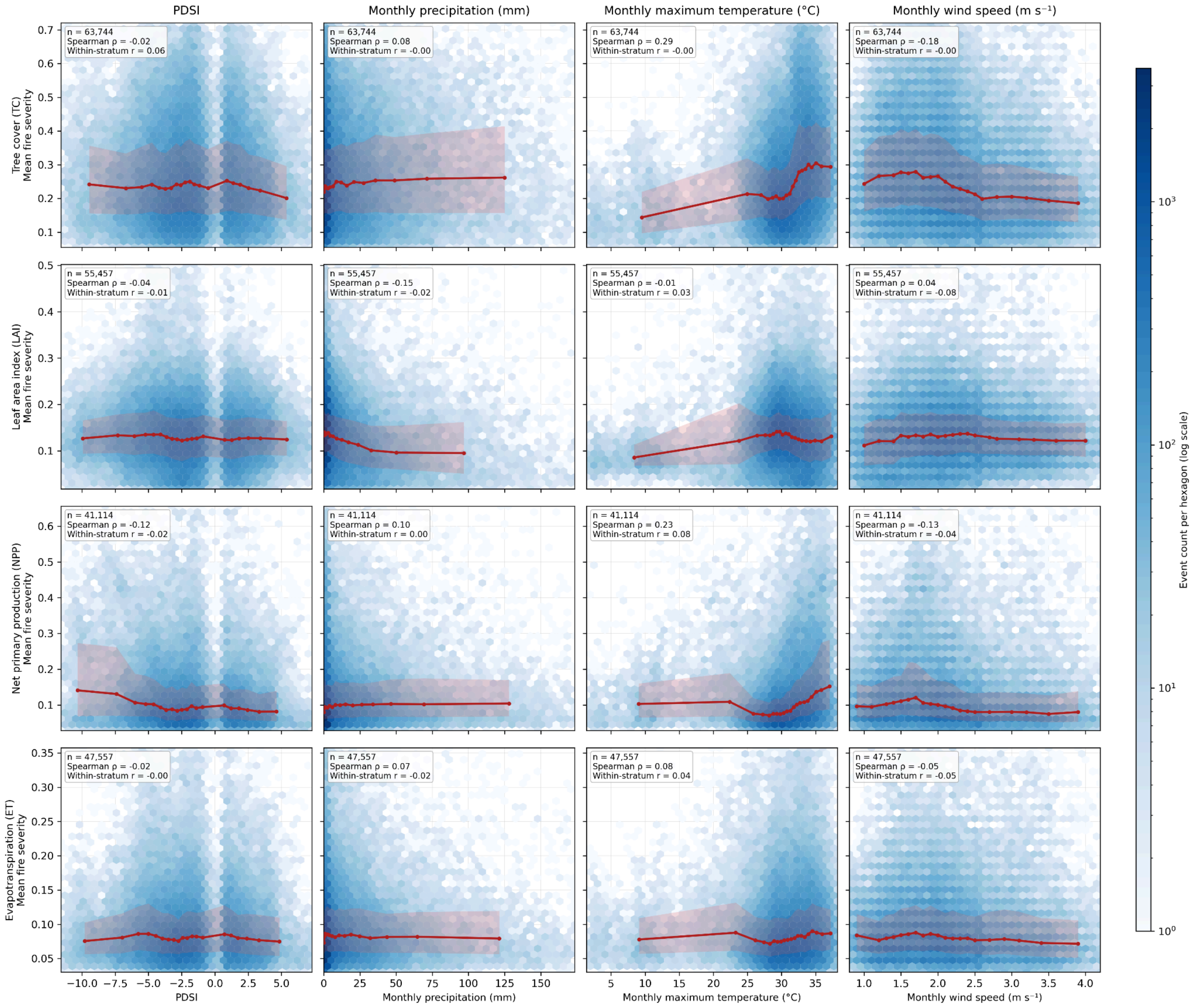


**Figure S21. Relationships between monthly fire-weather indicators and vegetation-based fire severity.** Rows show event-level mean fire severity derived from tree cover (TC), leaf area index (LAI), net primary productivity (NPP), and evapotranspiration (ET), while columns show PDSI, monthly precipitation, monthly maximum temperature, and monthly wind speed during the fire month. Blue hexagons show event density on a logarithmic count scale. Red curves represent median severity within equal-frequency weather bins, and pale red shading indicates the corresponding interquartile range. Insets report the number of eligible fire events, pooled Spearman correlation, and within-stratum rank association calculated within region × biome × fire-month strata. Panels use the metric-specific available sample, so event numbers differ among vegetation indicators. Overall, the weak and inconsistent pooled trends and near-zero within-stratum associations indicate no clear global relationship between monthly fire-weather conditions and vegetation-based fire severity.
